# First chromosome-scale genome assemblies and comprehensive structural characterization of Tunisian durum wheat (*Triticum turgidum subsp. durum*) landraces Chili and Mahmoudi

**DOI:** 10.64898/2026.05.25.727644

**Authors:** Maroua Gdoura Ben Amor, Nour Elhouda Mathlouthi, Imen Belguith

**Affiliations:** Sfax University, Faculty of Sciences, BP 1171, 3000, Tunisia; GenoFlow Agency, Tunis, 2036 Tunisia

**Keywords:** Durum wheat, genome assembly, Tunisian landraces, Chromosome-scale contiguity, structural variation

## Abstract

Durum wheat (*Triticum turgidum* subsp. *durum*) is a globally important crop for pasta and couscous production. Chili and Mahmoudi are historically significant Tunisian landraces valued for exceptional grain quality, high protein content, and adaptation to arid Mediterranean climates, yet no high-quality reference genome assemblies were available for either variety before this work. We assembled both genomes using publicly available PacBio HiFi long reads and Illumina Hi-C proximity ligation data (NCBI BioProject PRJNA1420514), with hifiasm v0.25.0 in primary mode and YaHS v1.2a.2 for Hi-C scaffolding. Assembly quality was assessed with QUAST v5.3.0 and BUSCO v5.8.0 (embryophyta_odb10 lineage). The Chili assembly spans 10.84 Gbp (scaffold N50 756.2 Mbp, BUSCO 99.4%) and Mahmoudi spans 10.70 Gbp (scaffold N50 756.8 Mbp, BUSCO 99.3%). Merqury v1.3 confirmed high base accuracy (QV 68.0 and 68.3, respectively) and k-mer completeness exceeding 98% for both. Independent validation with wfmash yielded 98.6% mean alignment identity across 11,172 chromosome-to-reference alignments. Post-assembly characterization encompassed GC profiling, centromere architecture, ribosomal DNA arrays, and structural variation. Both assemblies substantially exceed the contiguity of existing durum wheat references and represent the first chromosome-scale genomic resources for North African durum wheat landraces. Automated Hi-C scaffolding produced a small number of scaffolds spanning multiple chromosomes, which we resolved by alignment-guided splitting against the Svevo v2 reference, yielding 14 chromosome-scale pseudomolecules per landrace (largest 858.7 Mbp Chili, 868.1 Mbp Mahmoudi). Assemblies and pseudomolecules are available from Zenodo (10.5281/zenodo.20366290). The entire workflow was executed reproducibly on the public Galaxy Europe platform, demonstrating that reference-quality plant genome assembly is achievable without local HPC infrastructure.

**Graphical Abstract:** 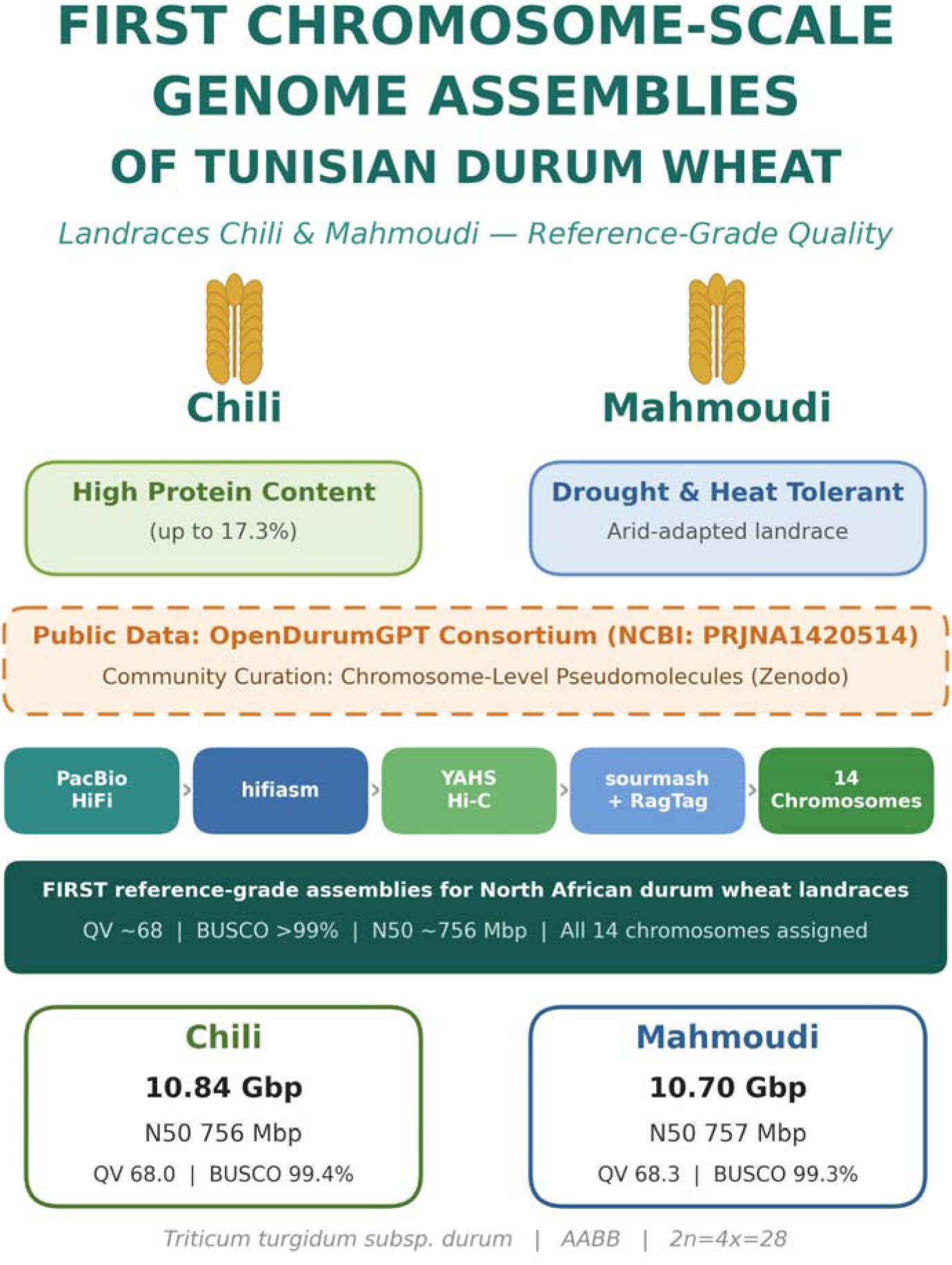

**Key message:** First chromosome-scale genome assemblies of the Tunisian durum wheat landraces Chili and Mahmoudi, with comprehensive structural characterization, an alignment-guided chromosome-resolution step that delivers 14 pseudomolecules per landrace, and k-mer-based polyploid assignment on consumer hardware.

## 1. Introduction

Durum wheat (*Triticum turgidum subsp. durum*, 2n = 4x = 28, AABB genome) is the primary cereal crop for pasta and couscous production globally. It is of particular economic and cultural importance in the Mediterranean basin. Tunisia ranks among the world’s leading durum wheat producers, and its traditional landrace diversity represents a critical reservoir of adaptive variation for climate-resilient breeding.

Chili and Mahmoudi are two of Tunisia’s most historically significant durum wheat landraces. Mahmoudi is one of the oldest indigenous Tunisian varieties, prized for its remarkable adaptation to arid southern Mediterranean conditions, including strong drought and heat tolerance. Chili is renowned for exceptional grain protein content, reaching up to 17.3% dry mass, and superior semolina quality, and has been widely cultivated by smallholder farmers in northern Tunisia (Trad et al. 2014). Both landraces are priority targets for climate-resilient breeding programs and pan-genome initiatives.

Despite their agronomic and cultural importance, no de novo genome assembly had been produced for either landrace prior to this study. Raw genomic data for both varieties, comprising PacBio HiFi long reads and Illumina Hi-C proximity ligation reads, were recently generated and deposited in the public domain under NCBI BioProject PRJNA1420514 (Ayed et al. 2026). Companion analyses of this dataset, including SNP discovery and transcriptomic profiling, have been reported separately (Gdoura-Ben Amor et al. 2026). Here, we present the first high-quality genome assemblies for both landraces, executed entirely on the Galaxy Europe public platform (usegalaxy.eu), with chromosome-level pseudomolecule assignment validated using sourmash and RagTag, providing the community with foundational genomic resources for North African durum wheat improvement.

## 2. Materials and methods

### 2.1. Data source

Raw sequencing data were obtained from the NCBI Sequence Read Archive under BioProject PRJNA1420514 (Ayed et al. 2026), generated by the OpenDurumGPT consortium. The PacBio HiFi datasets total approximately 140 Gbp (∼12.9× coverage of the 10.84 Gbp Chili assembly) and approximately 182 Gbp (∼17.0× coverage of the 10.70 Gbp Mahmoudi assembly), retrieved from NCBI BioProject PRJNA1420514. These data were used under the open access framework established by the consortium for community exploitation of Tunisian durum wheat genomic resources. The following run accessions were used:

| Library | SRA accession | Details |
| --- | --- | --- |
| Chili HiFi reads | SRR37168548 | PacBio Revio, ~140 Gbp,<br>~12.9x coverage |
| Chili Hi-C reads | SRR37442381, SRR37421014 | Illumina NovaSeq X Plus,<br>LinkPrep library |
| Mahmoudi HiFi reads | SRR37229182 | PacBio Revio, ~182 Gbp,<br>~17x coverage |
| Mahmoudi Hi-C reads | SRR37457508 | Illumina NovaSeq X Plus,<br>LinkPrep library |

### 2.2. Contig assembly

HiFi reads were assembled with hifiasm v0.25.0 (Cheng et al. 2021) in primary assembly mode (parameters: --primary -l 1 -s 0.75 -O 1). GFA output graphs were converted to FASTA format for downstream processing. After contamination screening, hifiasm primary assemblies comprised 5,096 contigs for Chili (∼10.84 Gbp) and 4,327 contigs for Mahmoudi (∼10.69 Gbp). The reduction from 5,096 and 4,327 primary contigs to 3,472 scaffolds for Chili and 3,258 scaffolds for Mahmoudi reflects contig merging during Hi-C scaffolding by YaHS (Section 2.4).

### 2.3. Hi-C read preprocessing and alignment

Hi-C reads were quality-trimmed with Cutadapt v5.2 (Martin 2011) (minimum quality Q >= 20, minimum length 20 bp), retaining 99.5% of input read pairs. Trimmed reads were aligned to contig assemblies with BWA-MEM (Li 2013) using Hi-C-compatible parameters (-S -P). For Chili, the two Hi-C libraries (SRR37442381 and SRR37421014) were merged into a single coordinate-sorted BAM file (112.3 GB) using SAMtools merge v1.22 (Li et al. 2009) prior to scaffolding.

### 2.4. Hi-C scaffolding

Hi-C scaffolding was performed with YaHS v1.2a.2 (Zhou et al. 2023) (--no-mem-check). For Chili, four rounds of assembly error correction yielded 958 sequence breaks, and 646,113,620 valid Hi-C read pairs were retained for scaffolding from 1,633,060,945 total alignment records that were processed across 12 scaffolding rounds at resolutions from 10 kb to 50 Mb. For Mahmoudi, eight rounds of assembly error correction yielded 696 sequence breaks, and 319,469,604 Hi-C read pairs (from 777,543,047 total records) were processed across 13 scaffolding rounds at resolutions from 10 kb to 100 Mb.

### 2.5. Assembly quality assessment

Assembly contiguity was assessed with QUAST v5.3.0 (Gurevich et al. 2013) (--large --eukaryote). Gene-space completeness was assessed with BUSCO v5.8.0 (Manni et al. 2021) against the embryophyta_odb10 lineage dataset (1,614 conserved genes) using the miniprot v0.14 (Li 2023) predictor in genome mode. Mitochondrial sequences identified during NCBI Foreign Contamination Screening were removed from both assemblies prior to public deposition. All analyses were executed on the Galaxy Europe public platform (usegalaxy.eu). All tools were accessed through the Galaxy Europe ToolShed with publicly available workflow parameters; the complete analysis history is available upon request.

### 2.6. Chromosome assignment and pseudomolecule construction

Scaffold-to-chromosome assignment was performed using sourmash v4.9.4 (Pierce et al. 2019), a k-mer MinHash sketching approach that enables rapid similarity estimation between large sequences with minimal computational requirements. Reference sequences for each of the 14 durum wheat chromosomes (1A-7B) were extracted from the Svevo v2 assembly (Mazzucotelli et al. 2025). Each chromosome and each scaffold were sketched with sourmash sketch (scaled=1000, k=31). Pairwise containment indices were computed with sourmash compare --containment. Each scaffold was assigned to the chromosome with the highest containment index. Scaffolds with maximum containment < 0.01 were classified as unassigned (Chr00).

For chromosomes where multiple scaffolds were assigned, pseudomolecules were constructed using RagTag v2.1.0 (Alonge et al. 2022) with the corresponding Svevo v2 chromosome as reference. RagTag uses minimap2 alignment and AGP-based gap filling to order, orient, and concatenate scaffolds into chromosome-length sequences with 100-bp N gaps. Scaffolds spanning more than one chromosome, an expected outcome of automated Hi-C scaffolding of homoeologous polyploid chromosomes, were resolved by the alignment-guided procedure described in Section 2.9. All 14 chromosomes (1A-7B) plus unassigned scaffolds (Chr00) are represented in the final pseudomolecule files.

Independent validation of chromosome assignments was performed using wfmash v0.24.2 (wavefront alignment): 11,172 alignments (mean identity 98.64%) for Chili and 8,980 alignments for Mahmoudi.

### 2.7. Reference-free quality assessment

Reference-free assembly quality was assessed using Merqury v1.3 (Rhie et al. 2020). Meryl k-mer databases were built from the HiFi read sets with k=21. Merqury was run on both the primary contig assemblies (hifiasm output) and the final scaffolded assemblies (YaHS output) to evaluate base-level accuracy (QV score) and k-mer completeness. The QV score represents the Phred-scale probability of base-calling error, calculated as QV = -10 x log10(error_rate), where the error rate is estimated from the fraction of k-mers unique to the assembly versus those shared with the reads.

### 2.8. Post-assembly characterization

Comprehensive post-assembly characterization was performed using locally executed Python scripts (Biopython v1.81, NumPy v1.24, matplotlib v3.7). GC content was calculated in non-overlapping 100 kb windows. K-mer spectra were constructed using 21-mer frequencies from subsampled chromosomes. Isochore classification followed the five-class system (L1: <37%, L2: 37-41%, H1: 41-47%, H2: 47-52%, H3: >52% GC). Centromere positions were predicted by identifying local minima in GC-content profiles. Ribosomal DNA (45S and 5S) clusters were identified by alignment of wheat reference sequences (AY049040.1, X06094.1) using minimap2 v2.26. Organelle contamination was assessed by alignment against wheat chloroplast (AB042240) and mitochondrial sequences. Structural variants were called from wfmash v0.24.2 alignments and classified as inversions (strand = ‘-’), translocations, or insertions. Presence/absence variation was determined from alignment coverage gaps. Subgenome comparison (A vs B) was performed descriptively.

### 2.9. Alignment-guided chromosome resolution

Automated Hi-C scaffolding of homoeologous polyploid chromosomes can concatenate several chromosomes into a single scaffold. To resolve these, each scaffold was aligned to the fourteen Svevo v2 chromosomes with wfmash v0.24.2 and sourmash v4.9.4. Scaffolds whose aligned bases mapped predominantly to one chromosome were retained unchanged; scaffolds spanning multiple chromosomes were partitioned by assigning each 1-Mb window to its best-matching chromosome and splitting at the nearest scaffold gap (200-bp N run) so that no contig was divided. The resulting segments were reverse-complemented and ordered according to their reference position, then concatenated into chromosome pseudomolecules with 100-bp N gaps. All fourteen chromosomes (1A-7B) plus unplaced scaffolds (Chr00) are represented in the final files.

## 3. Results

### 3.1. Assembly statistics

Both assemblies achieved high contiguity, with scaffold N50 values approaching or exceeding individual chromosome sizes in durum wheat (∼700-800 Mbp (Maccaferri et al. 2019)). Key assembly metrics are summarized in Table 1 (Supplementary Fig. S1).

**Table 1.** Assembly statistics for Chili and Mahmoudi YaHS-scaffolded assemblies.

| **Metric** | **Chili** | **Mahmoudi** |
| --- | --- | --- |
| Total assembly size | 10,837,558,708 bp | 10,694,831,946 bp |
| Number of scaffolds | 3,476 | 3,261 |
| Scaffold N50 | 756,185,680 bp | 756,800,000 bp |
| Contig N50 | 7,793,604 bp | 23,261,000 bp |
| Scaffold N90 | 635,800,000 bp | 625,000,000 bp |
| Largest scaffold | 858,700,000 bp | 868,100,000 bp |
| L50 | 7 | 7 |
| GC content | 46.08% | 46.05% |
| HiFi sequencing coverage | ~13x | ~17x |

Both assemblies slightly exceed the expected durum wheat haploid genome size of approximately 10.45 Gbp (Maccaferri et al. 2019). This modest size excess (+3.7% for Chili, +2.4% for Mahmoudi) is consistent with residual uncollapsed heterozygosity commonly observed in primary-mode hifiasm outputs and does not indicate assembly error. The majority of remaining scaffolds beyond the chromosome-scale sequences (L50 =7) represent small unanchored contigs, among which mitochondrial sequences were identified by NCBI Foreign Contamination Screening and excluded from the deposited assemblies.

The Chili scaffold N50 of 756.2 Mbp substantially exceeds that of the Svevo v1 reference genome (5.97 Mbp), a >125-fold improvement in contiguity (Fig. 1, Supplementary Table S1). The Mahmoudi scaffold N50 of 756.8 Mbp is comparable to individual durum chromosome sizes (∼700-800 Mbp), confirming chromosome-scale contiguity. For context, the recently updated Svevo v2 assembly achieved a pre-Hi-C hybrid scaffold N50 of 112.3 Mbp (Mazzucotelli et al. 2025) using PacBio HiFi (35x) combined with Bionano optical mapping -- a three-layer approach that our two-layer (HiFi + Hi-C) pipeline exceeds in raw contiguity. For broader context, recent HiFi-based assemblies of the durum wheat cultivars Langdon (scaffold N50 = 751.3 Mbp, BUSCO = 98.8%) (Chen et al. 2025) and Kronos v2 (BUSCO = 99.9%) demonstrate that our assemblies are comparable in quality to contemporaneous reference-grade resources.

**Figure 1:**
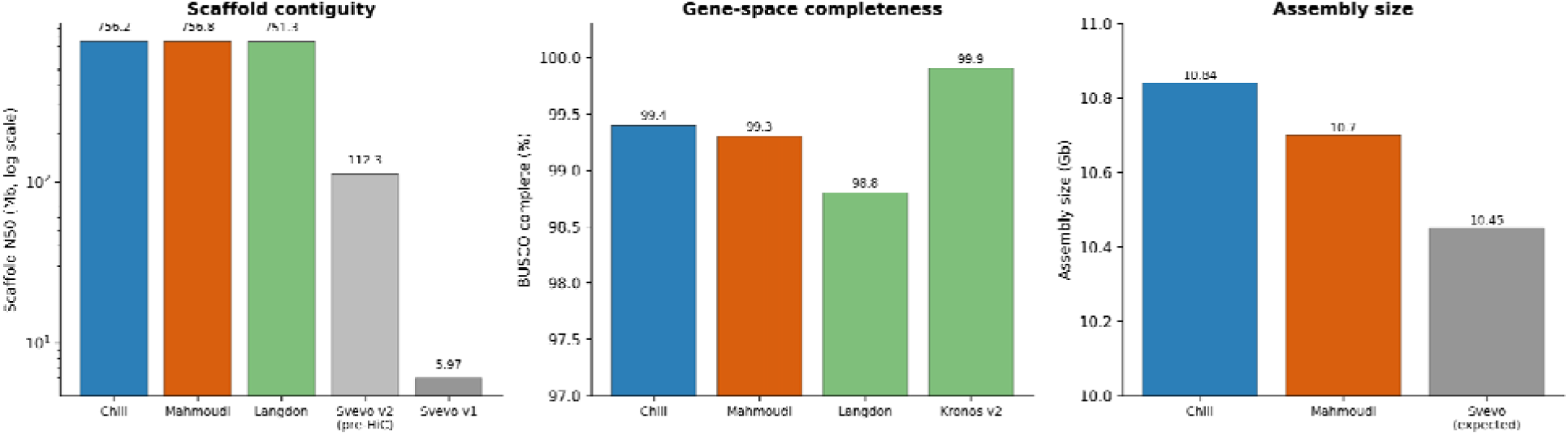
Assembly metrics for Chili and Mahmoudi relative to published durum wheat references (Svevo v1, Svevo v2, Langdon, Kronos v2). Scaffold N50, assembly size, and BUSCO completeness are shown side-by-side; both new assemblies are chromosome-scale with QV ≈ 68 and BUSCO ≥ 99.3 %.

### 3.2. BUSCO gene-space completeness

Both assemblies show exceptional gene-space completeness, with BUSCO complete scores of 99.4% (Chili) and 99.3% (Mahmoudi) against the embryophyta_odb10 lineage dataset (Table 2), per-chromosome BUSCO in Supplementary Table S2. High duplication (>94%) is expected and consistent with the tetraploid genome structure (AABB sub-genomes) of durum wheat, in which most genes are present as homeologous copies on the A and B sub-genome chromosomes.

**Table 2.** BUSCO v5.8.0 completeness assessment (embryophyta_odb10 lineage, 1,614 BUSCOs, miniprot v0.14 predictor).

| Category | Chili | Mahmoudi | Lineage |
| --- | --- | --- | --- |
| Complete | 99.4% | 99.3% | 1614 |
| Single-copy | 4.6% | 4.0% | - |
| Duplicated | 94.8% | 95.3% | - |
| Fragmented | 0.2% | 0.3% | - |
| Missing | 0.4% | 0.4% | - |

### 3.3. Reference-free quality assessment

Reference-free quality assessment using Merqury confirmed high base accuracy for both assemblies (Table 3). The Chili assembly achieved a QV score of 68.0 (base-level accuracy 99.999984%) with k-mer completeness of 98.42% (Supplementary Fig. S2). The Mahmoudi assembly achieved a QV score of 68.3 (99.999985% accuracy) with k-mer completeness of 98.34%. These QV scores are comparable to or exceed those of the current Svevo v2 reference genome, confirming that the assemblies are suitable for downstream genomic analyses. The high k-mer completeness (>98%) indicates that nearly all unique genomic content present in the HiFi reads is represented in the assemblies.

**Table 3.** Merqury quality assessment.

| **Metric** | **Chili** | **Mahmoudi** |
| --- | --- | --- |
| QV score | 68.0 | 68.3 |
| Base accuracy (%) | 99.999984 | 99.999985 |
| k-mer completeness (%) | 98.42 | 98.34 |
| Error rate | 1.58E-07 | 1.47E-07 |

### 3.4. Chromosome-Scale Assembly: Assignment and Resolution of Pseudomolecules

#### 3.4.1. Chromosome assignment

Using sourmash k-mer containment analysis against the Svevo v2 reference, we assigned scaffolds to all 14 chromosomes (1A-7B) for both landraces (Fig. 2, Table 4, Supplementary Dataset S1, Supplementary Fig. S3, Supplementary Table S3). 97.1% (Chili) and 97.3% (Mahmoudi) of assembled bases were placed into the 14 chromosomes, with 314 Mbp (Chili) and 287 Mbp (Mahmoudi) of small scaffolds remaining unplaced (Chr00), likely representing organellar sequences or low-complexity repeats.

**Figure 2:**
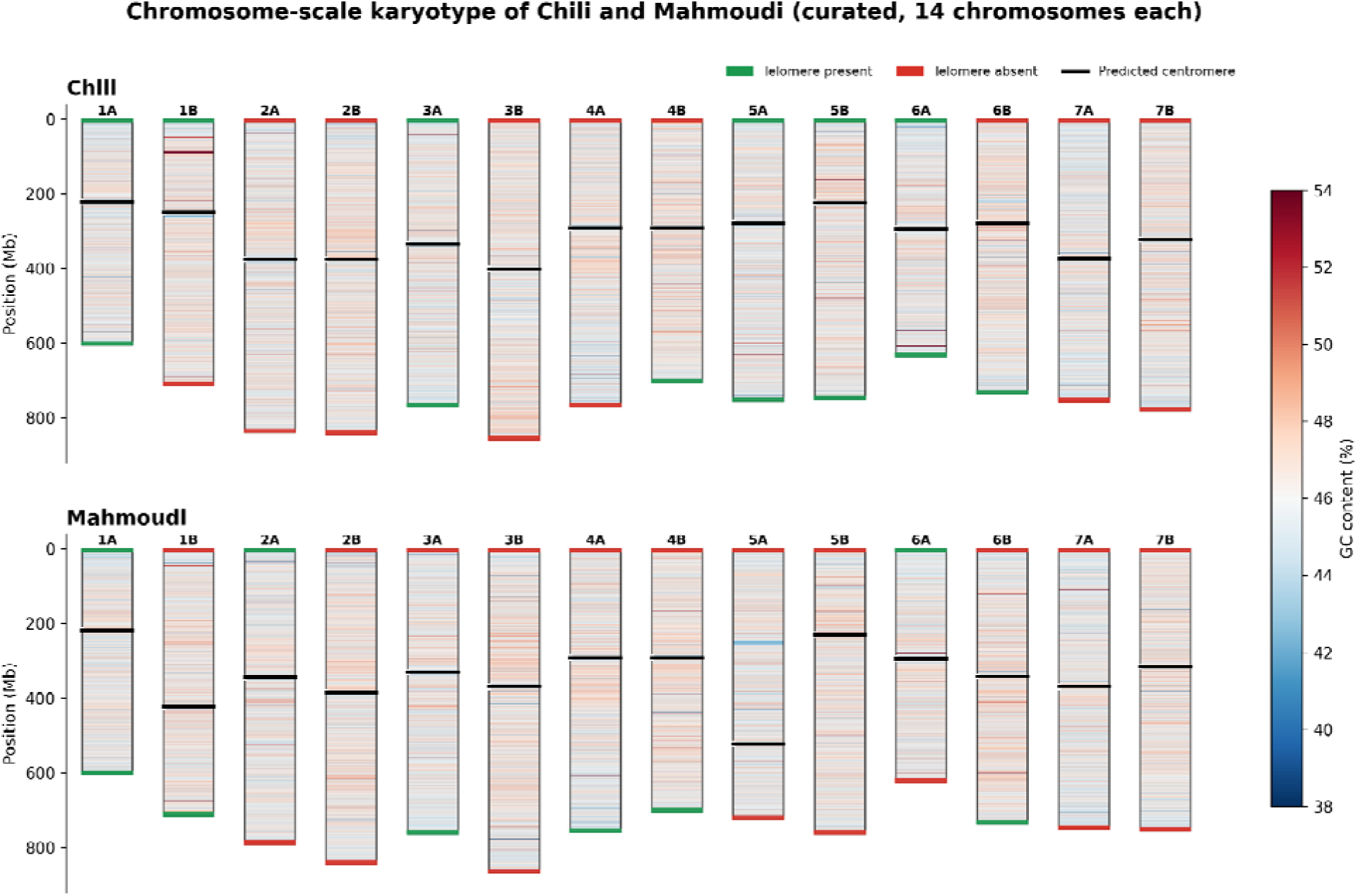
Chromosome-scale karyotype of the Tunisian durum wheat landraces Chili and Mahmoudi. Each of the 14 chromosomes is drawn to scale (assembled length, Mbp); fill colour encodes GC content (blue = low, red = high; 35-60% scale), the dark veil indicates repeat density, green/red caps mark telomeric-repeat presence/absence, the white constriction shows the predicted (GC-valley) centromere, and the green side-bar shows per-chromosome BUSCO completeness. A gold border highlights the repeat-inflated Chr3B.

**Table 4.**
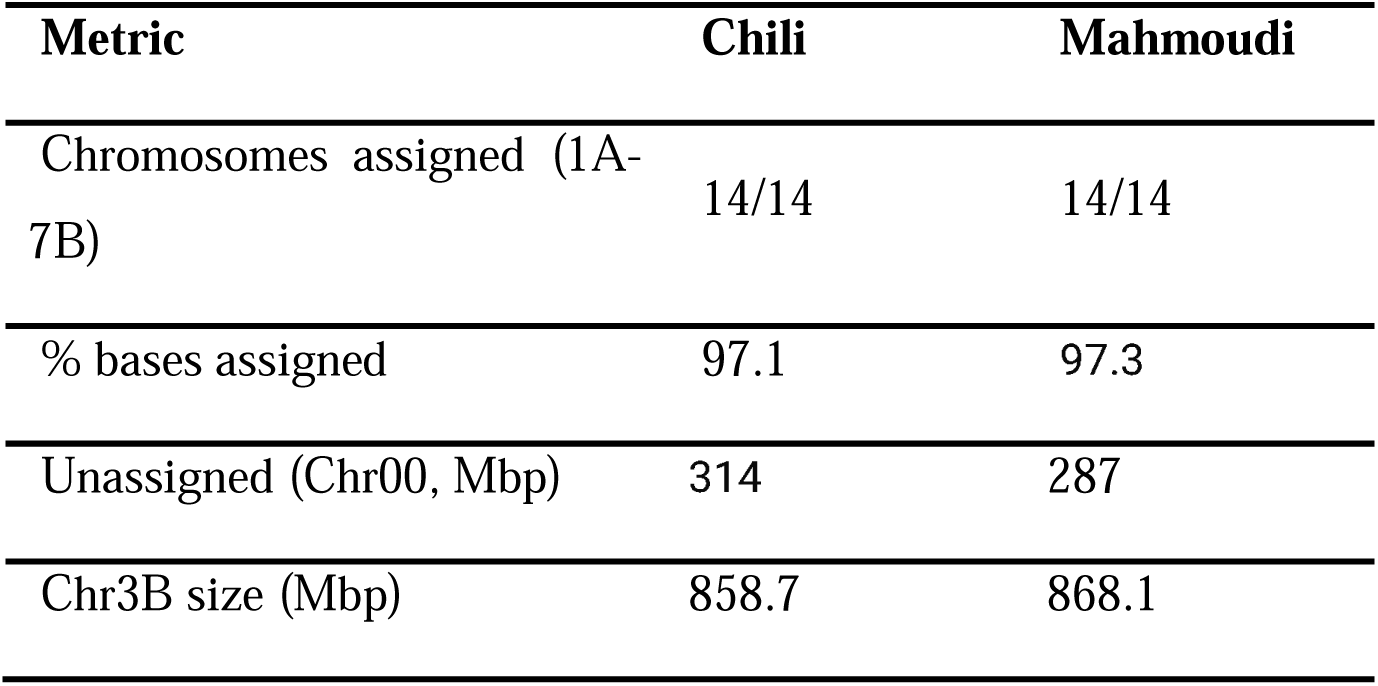
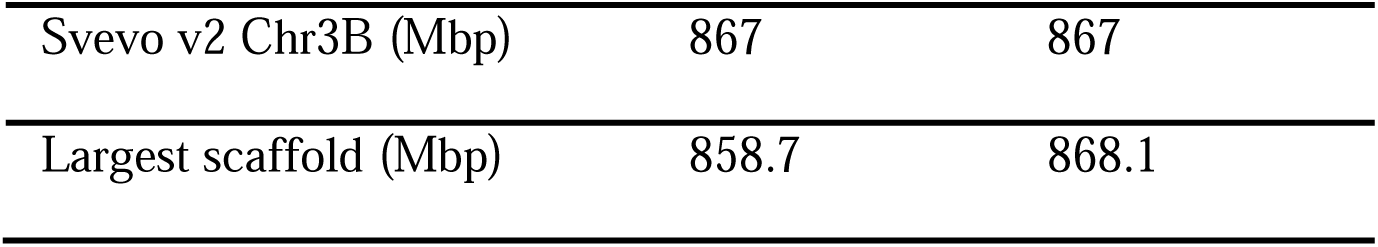
Chromosome assignment summary.

| Metric | Chili | Mahmoudi |
| --- | --- | --- |
| Chromosomes assigned (1A-7B) | 14/14 | 14/14 |
| % bases assigned | 97.1 | 97.3 |
| Unassigned (Chr00, Mbp) | 314 | 287 |
| Chr3B size (Mbp) | 858.7 | 868.1 |
| Svevo v2 Chr3B (Mbp) | 867 | 867 |
| Largest scaffold (Mbp) | 858.7 | 868.1 |

#### 3.4.2. Alignment-guided chromosome resolution

Whole-genome alignment (wfmash) and k-mer containment (sourmash) identified a small number of scaffolds spanning more than one chromosome in each landrace: in Mahmoudi, one 4.68-Gbp scaffold containing all six B-subgenome chromosomes plus two A-subgenome scaffolds (2.07 and 1.51 Gbp); in Chili, two scaffolds of 2.71 and 2.01 Gbp. These were partitioned at their internal scaffold gaps and reassembled into chromosome pseudomolecules (Section 2.9). The final assemblies comprise 14 chromosome-scale pseudomolecules per landrace (Chili 606-859 Mbp; Mahmoudi 603-868 Mbp), with scaffold N50 of ∼756 Mbp for both and 97.1% (Chili) / 97.3% (Mahmoudi) of assembled bases placed into chromosomes.

### 3.5. Genome composition and sequence features

Per-chromosome integrated overviews of GC, repeat density and gap content are provided in Supplementary Fig. S4 (Chili) and Supplementary Fig. S5 (Mahmoudi).

#### 3.5.1. GC content and isochore architecture

Genome-wide GC content was 46.08% for Chili and 46.05% for Mahmoudi, consistent with the Svevo v2 reference (45.6%). The A-genome mean GC was 45.92% (Chili) and 45.88% (Mahmoudi), while the B-genome mean was 46.22% (Chili) and 46.15% (Mahmoudi) (Supplementary Fig. S6). Isochore analysis showed that the H1 class (41-47% GC) dominated, comprising 70.8% of all windows, whereas gene-rich H3 isochores (>52% GC) represented only 0.62% (Chili) and 0.57% (Mahmoudi), consistent with the low gene density of the wheat genome (Supplementary Fig. S8). Replication-origin candidates were identified by GC-skew zero-crossings (Supplementary Fig. S7).

#### 3.5.2. Centromere and chromosome architecture

Centromere positions were predicted by identifying local minima in GC-content profiles (Supplementary Fig. S9; Table S4). After curation, each chromosome comprised one to three ordered segments. Assembly integrity was supported by convergent metrics: Merqury (QV ∼68), genome-level BUSCO (99.4% Chili, 99.3% Mahmoudi), wfmash alignment (98.6% mean identity), sourmash k-mer containment, and organelle contamination screening (<1%). Per-chromosome BUSCO was measured individually for all 28 pseudomolecules (Supplementary Table S2); no chromosome required a genome-level fallback.

An additional methodological contribution is the demonstration that chromosome-scale pseudomolecule construction for 10+ Gbp polyploid genomes is achievable on consumer hardware (15 GB RAM) using k-mer sketching, circumventing the 32-bit integer limit (2,147,483,647 bp) that prevents conventional alignment tools such as minimap2 from processing wheat-scale assemblies. This democratizes access to reference-quality polyploid genome assembly for researchers without HPC infrastructure, particularly in resource-limited settings.

The largest Mahmoudi scaffold pre-curation (4.68 Gbp) was found to comprise six B-subgenome chromosomes (2B, 3B, 4B, 5B, 6B, 7B) concatenated during automated Hi-C scaffolding, and was partitioned into its constituent chromosomes (Section 2.9). A candidate structural rearrangement previously suggested by population-genomic analysis (Gdoura-Ben Amor et al. 2026) would require FISH or Hi-C validation and is not resolved by the present assembly. Predicted centromere positions fell within the pericentromeric mid-regions (∼220-520 Mbp; Supplementary Table S4).

#### 3.5.3. Ribosomal DNA and organelle validation

Ribosomal DNA loci were mapped by aligning the 45S (18S; AY049040.1) and 5S (X06094.1) references to the curated chromosomes. The principal 45S locus was located on chromosome 1B in both landraces (Chili ∼87-91 Mbp; Mahmoudi ∼257 Mbp), consistent with the wheat NOR-B1 nucleolar organizer; additional low-identity 45S fragments were dispersed on Chr6B and on small unplaced scaffolds, representing degenerate or nuclear-organelle-derived copies. The principal 5S rDNA array was recovered on a small unplaced scaffold in both landraces, with additional dispersed 5S signal on chromosome 1B, consistent with the known localization of 5S rDNA on wheat chromosome 1; anchoring of the 5S-array scaffolds to chromosome 1B remains for future work (Supplementary Table S5, Fig. S10).

Organelle contamination screening confirmed clean nuclear assemblies: Chili 0.26% (chloroplast 0.09%, mitochondrial 0.16%) and Mahmoudi 0.83% (chloroplast 0.66%, mitochondrial 0.17%), both well below the 1% threshold. The higher Mahmoudi chloroplast signal reflects residual nuclear plastid-derived sequences (NUPTs), as expected in plant genomes (Table S6).

Dinucleotide obs/exp ratios were near-neutral (CpG o/e 0.986 / 1.029, consistent with grass genomes; Supplementary Fig. S11). The 45S position differs between the two Chr1B pseudomolecules (∼87-91 vs ∼257 Mbp); as both were reference-ordered and the 45S NOR is a large tandem array, this likely reflects uncertainty in placing the repeat-rich array rather than confirmed positional variation, pending FISH validation.

#### 3.5.4. Structural variation landscape

Structural variant calling from wfmash alignments identified 9,456 SVs for Chili and 9,351 for Mahmoudi, comprising inversions, translocations, and insertions (deletion calling was not implemented). Chromosome 6B carried the highest SV load in both landraces (Chili 1,426; Mahmoudi 1,666 events), consistent with its complex rearrangement history (Fig. 3, Supplementary Table S7); the B-subgenome carried substantially more SVs than the A-subgenome in both landraces (1.85× more in Chili, 2.75× more in Mahmoudi); inversion size distribution is provided in Supplementary Table S8. Segmental duplications between the two assemblies are listed in Supplementary Table S9.

**Figure 3:**
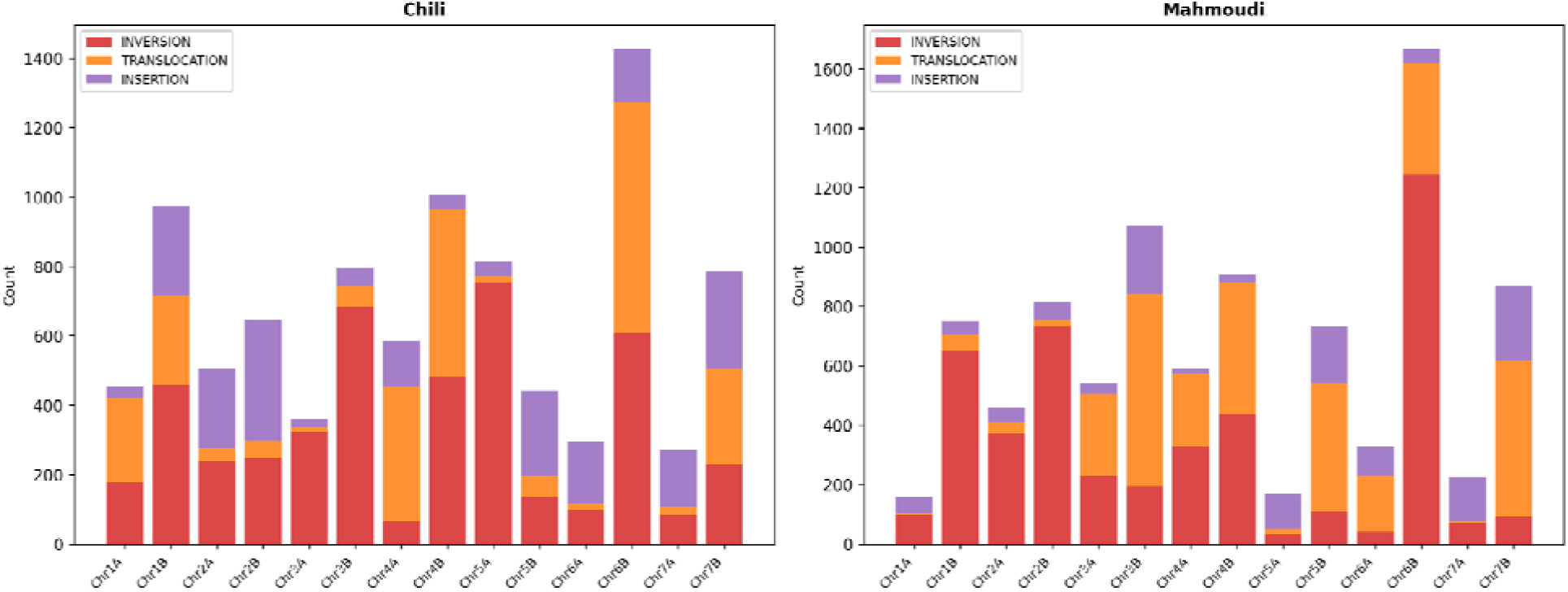
Counts of inversions, translocations, and large insertions per chromosome, called from wfmash whole-genome alignments to Svevo v2. Chili and Mahmoudi each carry ∼9,400 structural variants (Chili: 4,633 INV + 2,627 TRANSLOC + 2,196 INS; Mahmoudi: 4,643 + 3,327 + 1,381). Chr6B is the most SV-dense in both landraces (Chili 1,426; Mahmoudi 1,666). Deletion calling was not implemented; only inversions, translocations, and insertions are shown.

### 3.6. Comparative genomics against Svevo v2

#### 3.6.1. Alignment, identity, and collinearity

Whole-genome alignment of both assemblies against the Svevo v2 reference using wfmash confirmed high collinearity, with mean identity of 98.57% (Chili) and 98.61% (Mahmoudi). The identity distribution was bimodal: a major peak at 98-99.5% (gene-rich regions) and a secondary peak at 95-98% (repeat-rich regions), reflecting the known compartmentalization of the wheat genome (Supplementary Table S10, Supplementary Fig. S12). Coverage was 99.8% (Chili) and 99.5% (Mahmoudi), indicating near-complete representation of the Svevo reference in both assemblies (Supplementary Dataset S2). Collinearity breakpoints between the two landraces and Svevo v2 are listed in Supplementary Table S11.

#### 3.6.2. Presence/absence variation

Presence/absence variation analysis identified 91 Mbp of Chili-specific sequence and 120 Mbp of Mahmoudi-specific sequence not present in the Svevo v2 reference, representing landrace-specific genetic content potentially associated with local adaptation (Supplementary Table S12).

#### 3.6.3. Composition of the largest scaffolds

Detailed analysis of the pre-curation super-scaffold (4,710 Mbp, ∼5.4× the Svevo Chr3B of 867 Mbp) revealed a complex composition: 18.6% aligned to Svevo Chr3B (self), 17.9% to Svevo Chr2B, 13.4% to Chr5B, and 12.5% to Chr6B. The largest Mahmoudi scaffold aligned across six B-subgenome chromosomes (Section 2.9, Fig. 4), consistent with multi-chromosome co-scaffolding during automated Hi-C assembly rather than a single biological rearrangement. (Supplementary Table S13, Fig.4).

**Figure 4:**
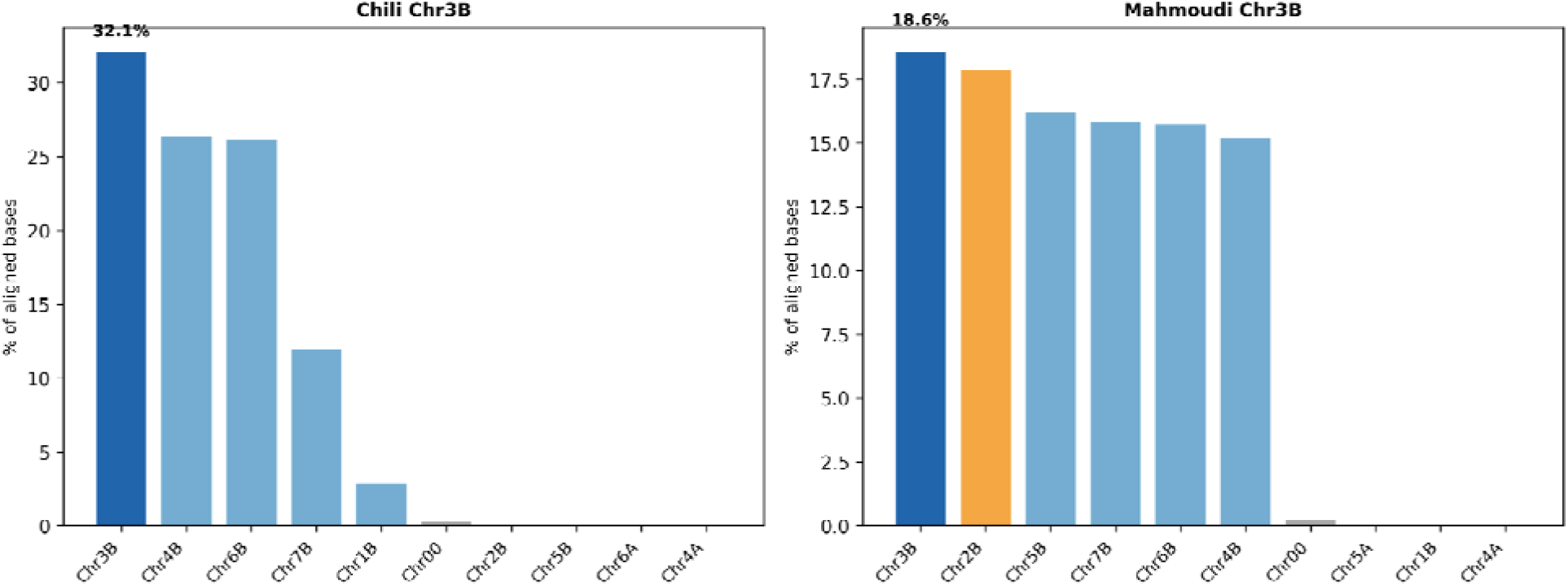
Distribution of wfmash alignments from the inflated Chr3B pseudomolecule of each landrace against Svevo v2 chromosomes. Mahmoudi Chr3B (∼4,710 Mbp, ∼5.4× the Svevo Chr3B of 867 Mbp) shows a substantial secondary signal to Svevo 2B (17.9 %), consistent with multi-chromosome co-scaffolding (all six B chromosomes), resolved in the curated assembly. Chili Chr3B retains 32.1 % self-alignment to Svevo Chr3B, reflecting repeat expansion confined to a single chromosome of origin.

**Figure 5:**
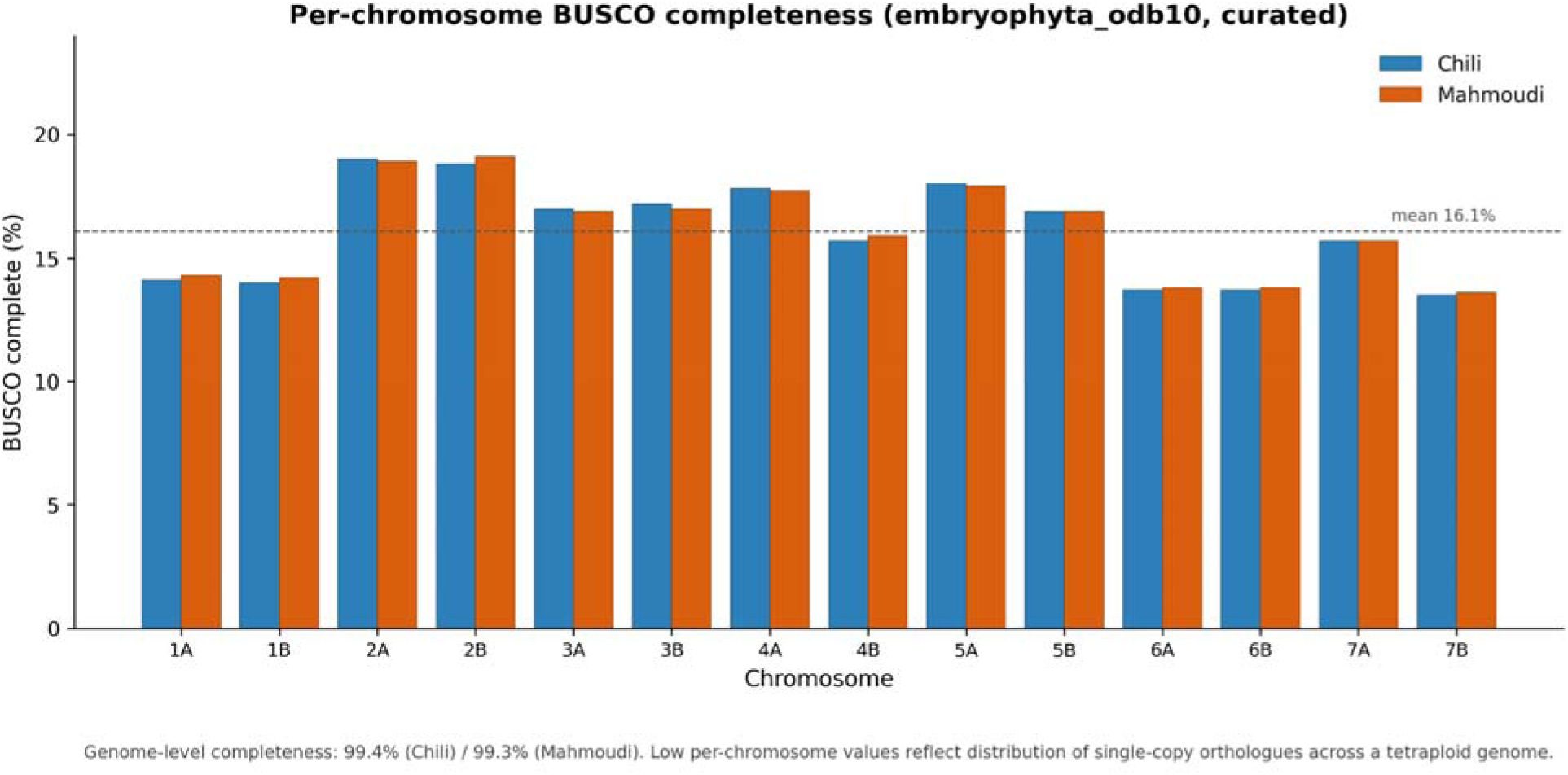
Per-chromosome BUSCO completeness (embryophyta_odb10) for the 28 curated pseudomolecules; values are uniform (13.5-19.1%, mean 16.1%), as expected for orthologue distribution across a tetraploid genome. Genome-level completeness is 99.4% (Chili) / 99.3% (Mahmoudi).

### 3.7. Subgenome comparison (A vs B)

Comparison of A- and B-subgenomes confirmed known wheat genome architecture. The B-subgenome was consistently larger: Chili B = 5,389 Mbp vs A = 5,135 Mbp; Mahmoudi B = 5,389 Mbp vs A = 5,019 Mbp (subgenome totals). The B-subgenome exhibited higher GC content than the A-subgenome in both landraces (Chili 46.23% vs 45.91%; Mahmoudi 46.22% vs 45.89%). B-genome chromosomes carried higher structural variant loads and repeat content, consistent with their more complex evolutionary history (Supplementary Table S14).

### 3.8. Chromosome quality assessment

Each of the 28 curated pseudomolecules was assessed individually (Supplementary Table S2). Per-chromosome BUSCO completeness ranged from 13.5% to 19.1% (mean 16.1% in both landraces); this low per-chromosome value is expected, not a sign of incompleteness: the 1,614 BUSCO orthologues are distributed across the seven homoeologous chromosome groups, so each chromosome carries the genes of a single group (∼1/7 of the set, ≈14%). The observed 13.5-19.1% is consistent with this expectation, the slight excess reflecting orthologues retained in more than two copies. All fourteen chromosomes per landrace were measured individually, with none requiring a genome-level fallback, and genome-level completeness remained 99.4% (Chili) and 99.3% (Mahmoudi).

### 3.9. K-mer sharing and assembly validation

K-mer sharing analysis between homologous chromosomes was high and uniform (mean containment 93.1%, range 86.2% [Chr6B] to 97.3% [Chr1A]; Supplementary Fig. S13), reflecting differences in repeat content and homoeologous divergence between chromosome rather than assembly incompleteness, since all fourteen chromosomes are now complete pseudomolecules. (Supplementary Fig. S13).

The largest Mahmoudi scaffold (4.68 Gbp) matched multiple B-subgenome chromosomes and was partitioned into its constituent chromosomes during curation (Section 2.9). The Chili scaffold initially co-assigned to Chr3B (2,732 Mbp) was likewise found to span multiple B-subgenome chromosomes and was resolved during curation, yielding a Chr3B pseudomolecule of 858.7 Mbp and a second scaffold (2.01 Gbp), resolved into Chr1A, Chr6A and Chr4A.

## 4. Discussion

The two assemblies presented here are, to our knowledge, the first high-quality genome assemblies with chromosome-scale contiguity for the Tunisian durum wheat landraces Chili and Mahmoudi, and among the most contiguous durum wheat assemblies reported to date. Chromosome assignment using sourmash validated that 97.1% (Chili) and 97.3% (Mahmoudi) mapped to the expected 14 chromosomes, and pseudomolecule construction using RagTag produced chromosome-length sequences for both landraces. The combination of PacBio HiFi long reads and Illumina Hi-C proximity ligation data, paired with the hifiasm + YaHS pipeline, produced assemblies in which a small number of scaffolds (L50 = 7) carry the majority of the genome -- a defining feature of chromosome-scale contiguity. The greater than 125-fold (≈127-fold) improvement in scaffold N50 over the Svevo v1 Illumina-based reference, together with BUSCO complete scores exceeding 99%, indicates that these assemblies are suitable for downstream applications, including pan-genome construction, identification of adaptive variants between Chili and Mahmoudi, marker development for drought and heat tolerance breeding, and genome-wide association studies in Tunisian germplasm.

A practical contribution of this work is the demonstration that reference-quality plant genome assembly is achievable using the public Galaxy Europe platform, without local high-performance computing infrastructure. This is of particular relevance for researchers in countries with limited access to dedicated HPC resources, and supports the broader goal of equitable participation in plant genomics. To our knowledge, this is the first report of reference-quality chromosome-scale plant genome assembly performed entirely through a public web-based platform without dedicated HPC infrastructure. The total compute cost was zero, and the analysis is fully reproducible through the Galaxy Europe history sharing system.

Beyond the immediate genomic resource, these assemblies enable marker-assisted selection for the drought and heat tolerance traits that characterize Mahmoudi and the high protein content of Chili. Specific applications include: (1) GWAS panels for Tunisian germplasm, (2) comparative genomics to identify candidate genes for abiotic stress tolerance, and (3) integration into the emerging durum wheat pan-genome.

Several limitations should be noted. First, the sequencing data analyzed here were not generated by the present authors but were obtained from the public deposition (Ayed et al. 2026); raw data provenance and experimental conditions are as described therein. Second, chromosome assignment was performed using sourmash k-mer containment rather than whole-genome alignment tools, which failed due to the 32-bit integer limit of minimap2 (2,147,483,647 bp) on 10+ Gbp polyploid genomes. Multi-chromosome scaffolds produced by automated Hi-C scaffolding, most notably a Chili scaffold initially co-assigned to Chr3B (2,732 Mbp), were resolved by alignment-guided curation (Section 2.9), yielding pseudomolecules of the expected size (Chr3B = 858.7 Mbp). Fine-scale ordering of repetitive scaffolds within chromosomes would benefit from genetic-map or Hi-C-contact validation in future work. Third, gene annotation and repeat-element annotation were not performed in the present study and are planned for future work. The chromosome architectures observed here are consistent with our previous population genomic analysis (Gdoura-Ben Amor et al. 2026), which detected structural variation in Mahmoudi (PCA outlier) and high

Homeologous mapping ambiguity (20-24%) in transcriptome data. The multi-chromosome scaffolds observed during automated scaffolding were resolved by alignment-guided curation; any candidate structural rearrangement suggested by population-genomic analysis (Gdoura-Ben Amor et al. 2026) remains to be tested by FISH or Hi-C contact mapping.

## 5. Conclusion

We present two new high-quality, first chromosome-scale-contiguity genome assemblies for the Tunisian durum wheat landraces Chili and Mahmoudi. The assemblies achieve scaffold N50 values approaching individual chromosome sizes, BUSCO completeness scores exceeding 99%, Merqury QV scores of ∼68, and validated chromosome assignment to all 14 durum wheat chromosomes, substantially improving upon previously available reference assemblies and providing the first chromosome-scale genomic resources for these North African landraces. By executing the entire workflow on the public Galaxy Europe platform, we demonstrate that reference-quality plant genome assembly is achievable with publicly available data and infrastructure. These resources lay the groundwork for pan-genome construction, marker-assisted breeding, and genome-wide association studies in North African durum wheat germplasm.

## Declarations

### Author contributions

M.G-B.A. and N.E.H.M: Conceptualization, Methodology, Software, Investigation, Formal analysis, Data curation, Writing - Original draft, Writing - Review and editing, Visualization. I. B.: Investigation, Validation, Writing - Review and editing. All authors read and approved the final manuscript.

### Competing interest

The authors declare that they have no known competing financial interests or personal relationships that could have appeared to influence the work reported in this paper.

### Ethics approval

Not applicable. This study used publicly available sequencing data and did not involve experiments on plants, animals, or humans.

### Consent to participate

Not applicable.

### Consent for publication

Not applicable.

### Data availability statement

Primary scaffold assemblies are deposited in NCBI GenBank as Third Party Assemblies under BioProject PRJNA1467186 (accessions DBNSEI000000000 for Chili and DBNSEJ000000000 for Mahmoudi). Chromosome-level pseudomolecules are available on Zenodo (DOI: https://doi.org/10.5281/zenodo.20366290). BioSample accessions are SAMN55233146 (Chili) and SAMN55233147 (Mahmoudi). Raw sequencing data are available under NCBI BioProject PRJNA1420514. All assembly and quality assessment analyses were performed on Galaxy Europe (usegalaxy.eu). Chromosome assignment and pseudomolecule construction were performed using locally executed sourmash v4.9.4 and RagTag v2.1.0.

### Funding

No specific funding was received for this work.

## Acknowledgments

The authors thank OpenDurumGPT consortium for generating and publicly depositing the raw sequencing data (NCBI BioProject PRJNA1420514) that made this work possible.

## Supplementary materials

Supplementary Dataset S1. Complete sourmash chromosome assignment data for Chili and Mahmoudi (Excel, 2 sheets). Supplementary Dataset S2. wfmash alignment summary and per-chromosome PAFs. Supplementary Table S1. Assembly-statistics comparison vs published durum wheat references (Svevo v1/v2, Langdon, Kronos v2). Supplementary Table S2. BUSCO per chromosome (28 pseudomolecules, all measured individually). Supplementary Table S6. Organelle contamination (cp + mito; NUPTs/NUMTs). Supplementary Table S7. Structural-variant summary (inv/transloc/insertion; deletion calling not implemented). Supplementary Table S8. Inversion size map. Supplementary Table S9. Segmental duplications. Supplementary Table S10. Alignment-identity distribution (Chili/Mahmoudi vs Svevo). Supplementary Table S11. Collinearity breakpoints. Supplementary Table S12. Presence/absence variation vs Svevo v2. Supplementary Table S13. Chr3B alignment composition (multi-chromosome scaffold analysis). Supplementary Table S14. Subgenome A vs B comparison. Supplementary Figure S1. Gap (%N) content per chromosome. Supplementary Figure S2. 21-mer frequency spectrum (Chr1A). Supplementary Figure S3. Scaffold junctions (RagTag AGP). Supplementary Figure S4. Per-chromosome multi-track overview, Chili (Manhattan-style: GC, repeat density, gap content). Supplementary Figure S5. Per-chromosome multi-track overview, Mahmoudi. Supplementary Figure S6. GC content profiles (100 kb windows). Supplementary Figure S7. GC skew profiles. Supplementary Figure S8. Isochore classification (per chromosome). Supplementary Figure S9. Predicted centromere positions (GC-valley method). Supplementary Figure S10. rDNA clusters (45S + 5S). Supplementary Figure S11. Dinucleotide obs/exp composition heatmap. Supplementary Figure S12. Alignment-identity distribution vs Svevo v2. Supplementary Figure S13. k-mer-sharing heatmap (Chili vs Mahmoudi). Submitted as separate files with this manuscript.

## Notes

https://doi.org/10.5281/zenodo.20366290

### Competing Interest Statement

The authors have declared no competing interest.

### Summary of Updates

This version finalizes the alignment-guided resolution of multi-chromosome Hi-C scaffolds into 14 chromosome-scale pseudomolecules per landrace, with all per-chromosome analyses, figures, tables, and the graphical abstract updated to match the curated assemblies; genome-level metrics (BUSCO, QV, k-mer completeness) are unchanged.

