## Supplementary material for "First chromosome-scale genome assemblies and comprehensive structural characterization of Tunisian durum wheat (*Triticum turgidum subsp. durum*) landraces Chili and Mahmoudi": Supplementary_Figures_S1-S13.docx

**Supplementary Figure S1. Gap (%N) content per chromosome**

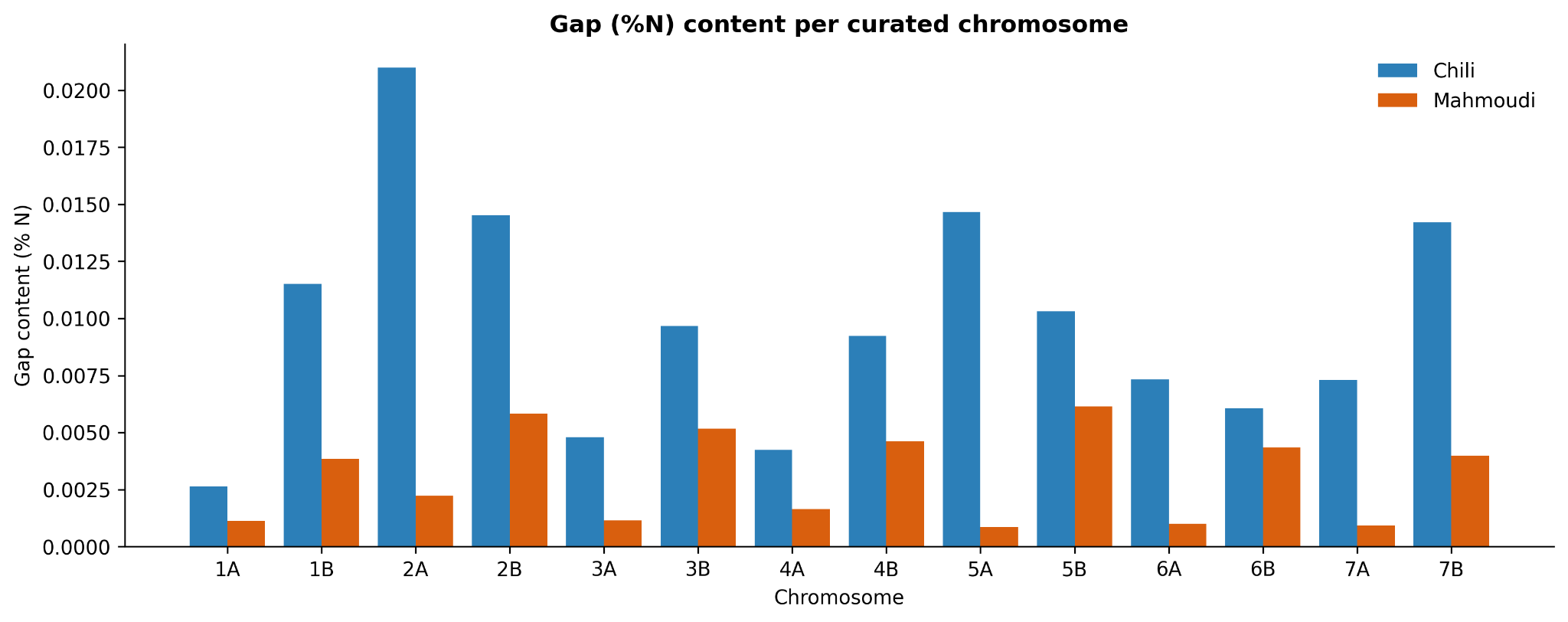

**Figure S1.** Bar chart of the percentage of N characters per pseudomolecule for Chili (green) and Mahmoudi (blue), and the corresponding number of N-blocks. Both assemblies show very low gap content (mean 0.050 % Chili, 0.058 % Mahmoudi), reflecting the high contiguity of the HiFi-based assemblies.

**Supplementary Figure S2. K-mer (21-mer) frequency spectrum, Chr1A**

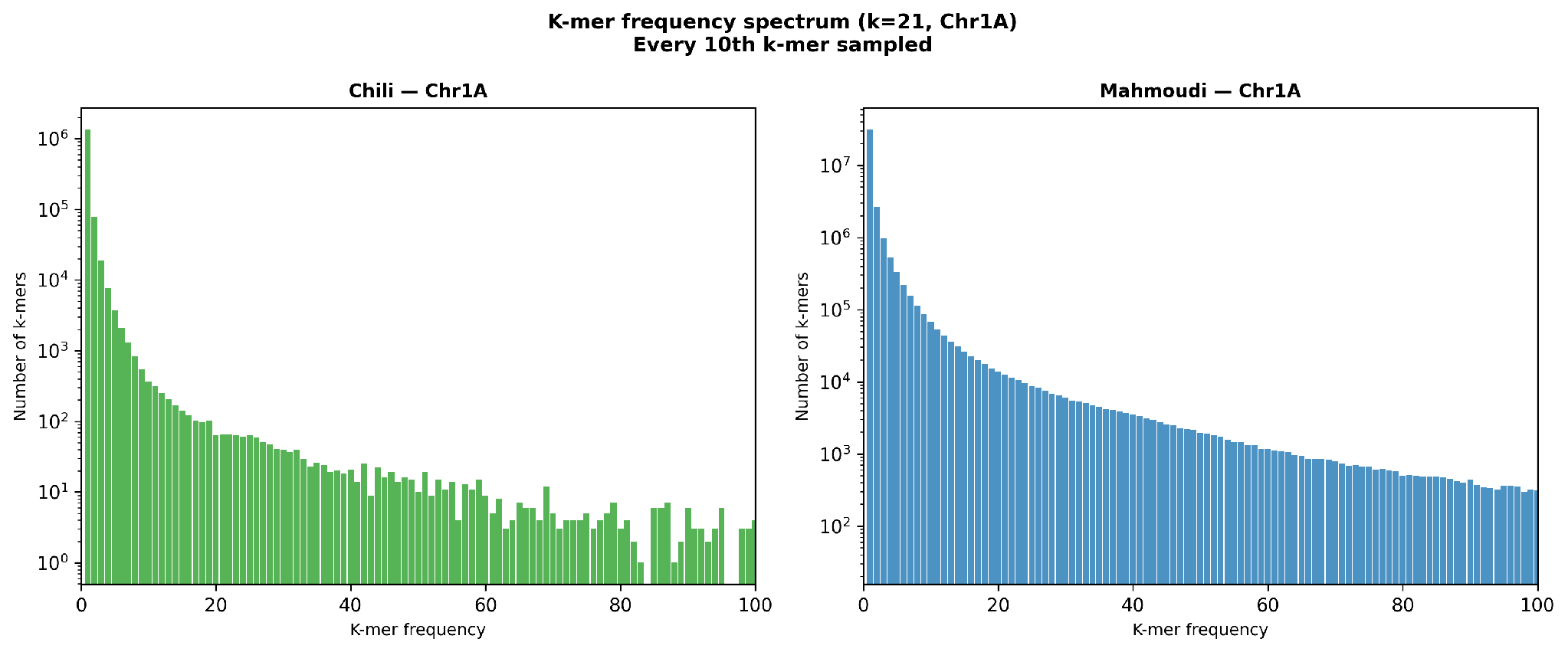

**Figure S2.** Frequency-of-frequency distribution of 21-mers along Chromosome 1A for Chili and Mahmoudi (every 10th 21-mer sampled). The spectrum shape is consistent with a tetraploid AABB genome; the 1x peak primarily reflects the sampling strategy.

**Supplementary Figure S3. Scaffold junctions per chromosome (RagTag AGP)**

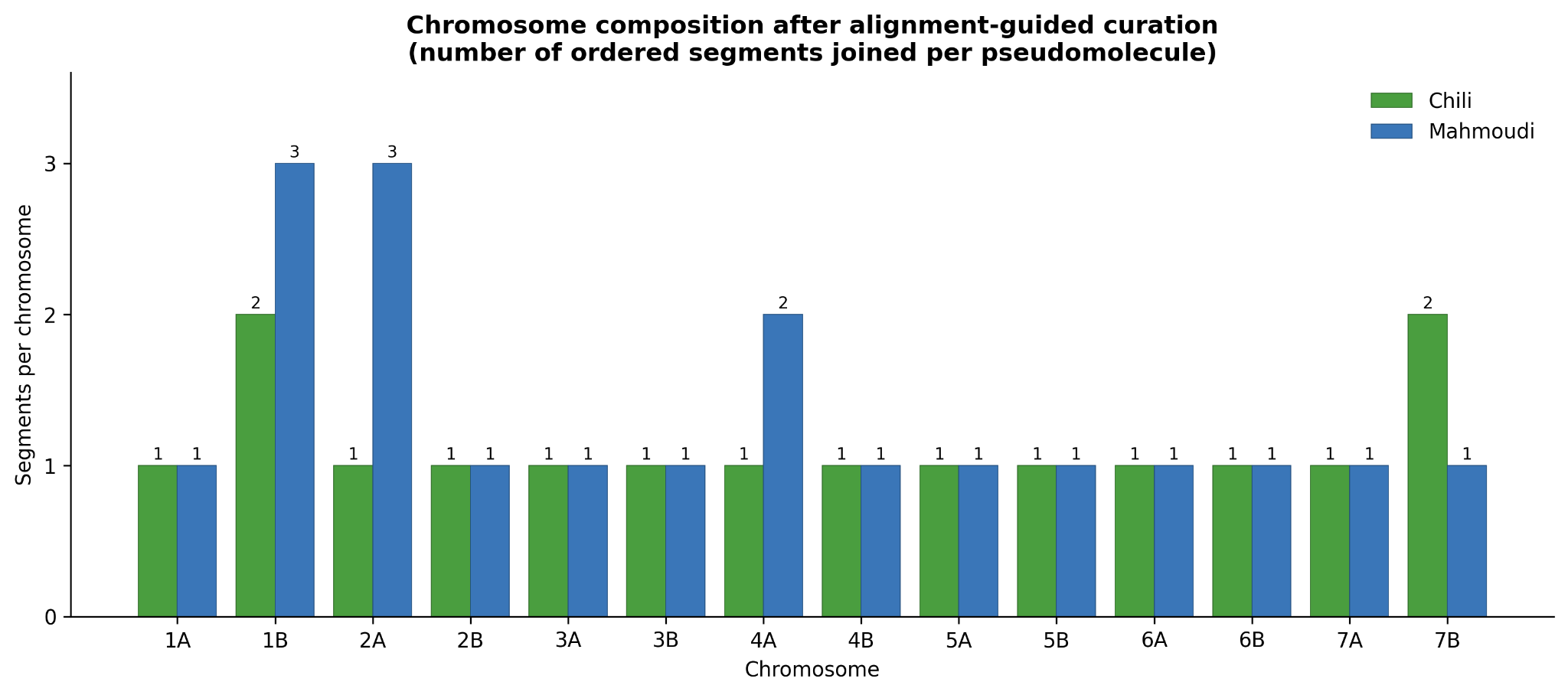

**Figure S3.** Chromosome composition after alignment-guided curation. Number of ordered segments joined to form each chromosome pseudomolecule for Chili (green) and Mahmoudi (blue). Most chromosomes comprise a single segment; a few were reassembled from multiple scaffolds ordered against the Svevo v2 reference (Mahmoudi Chr1B and Chr2A, three segments; Chr4A, two; Chili Chr1B and Chr7B, two).

**Supplementary Figure S4. Per-chromosome multi-track overview - Chili**

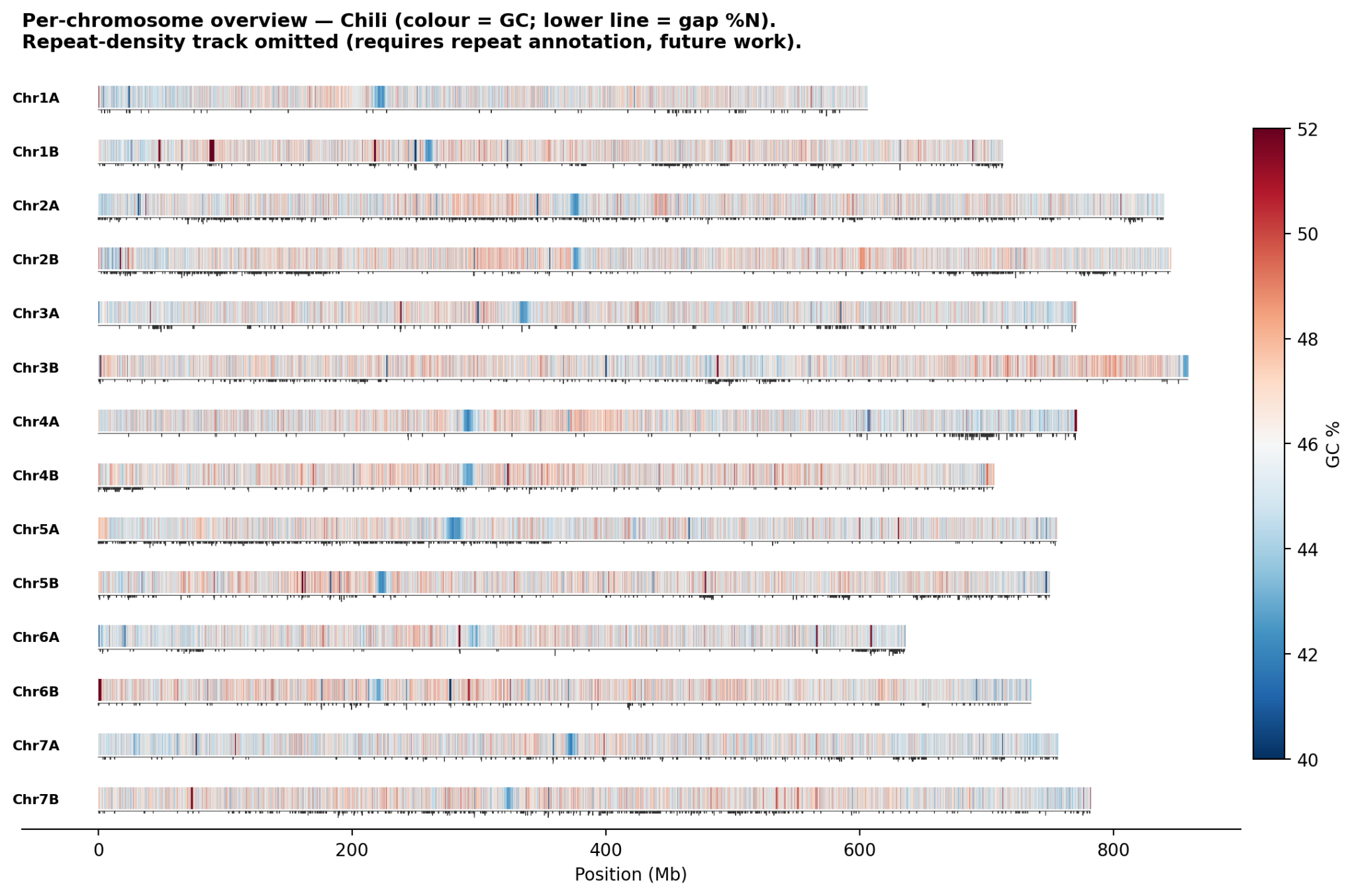

**Figure S4.** Per-chromosome summary for Chili: GC content (colour) and gap (%N) (lower line); repeat-density track omitted (requires repeat annotation, future work).

**Supplementary Figure S5. Per-chromosome multi-track overview - Mahmoudi**

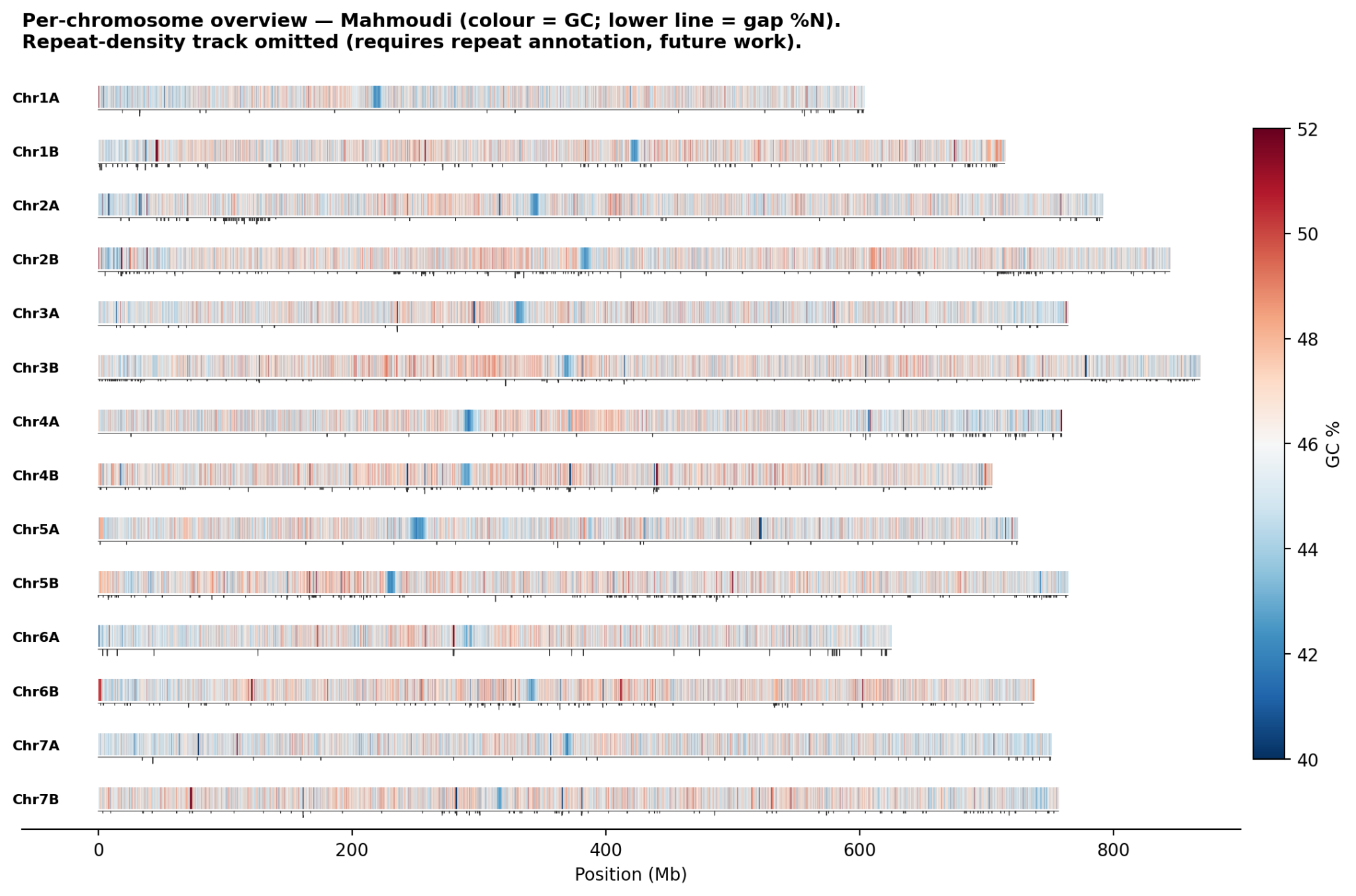

**Figure S5.** Per-chromosome summary for Mahmoudi: GC content (colour) and gap (%N) (lower line); repeat-density track omitted (requires repeat annotation, future work).

**Supplementary Figure S6. GC content profiles (100 kb windows)**

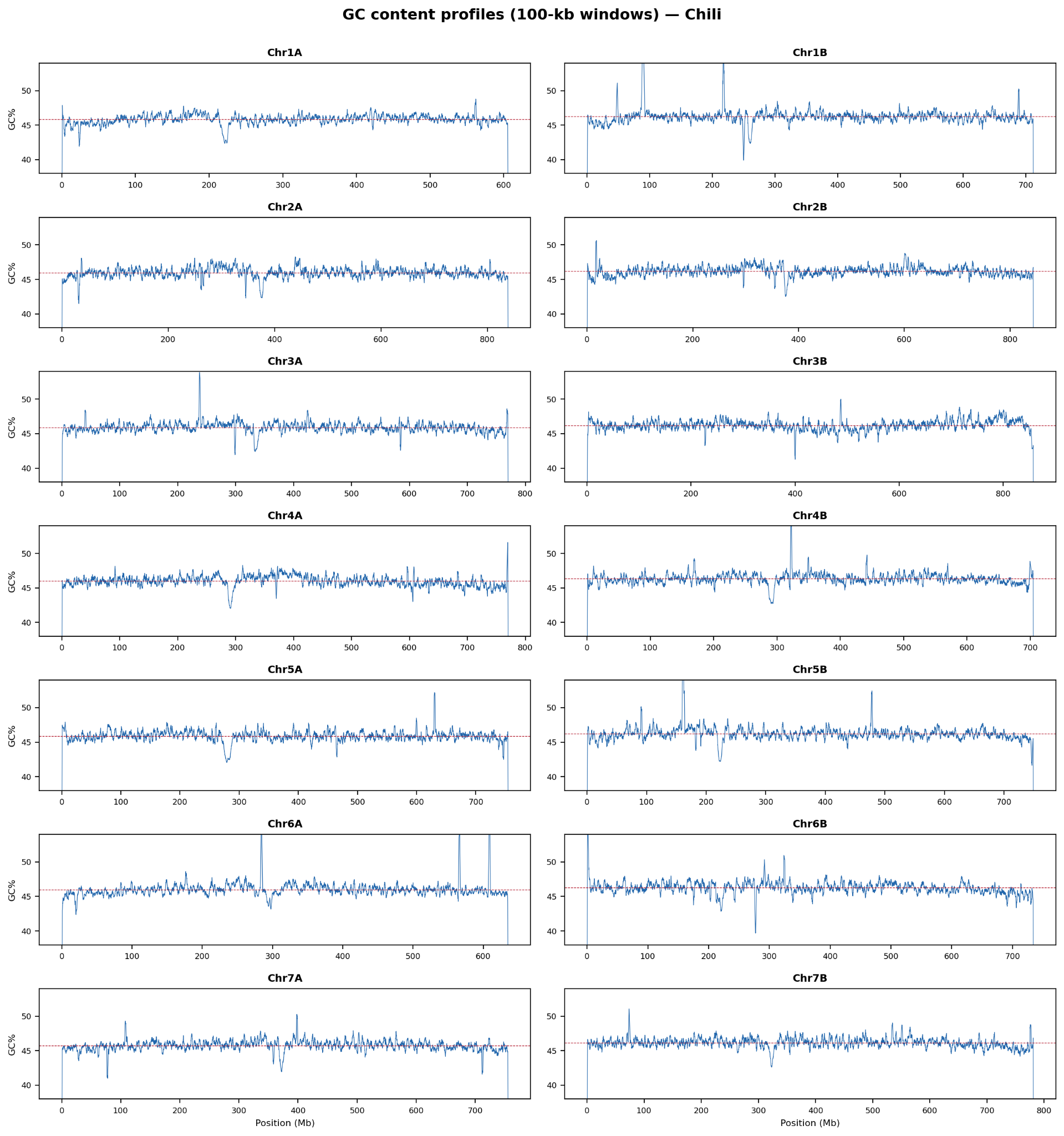

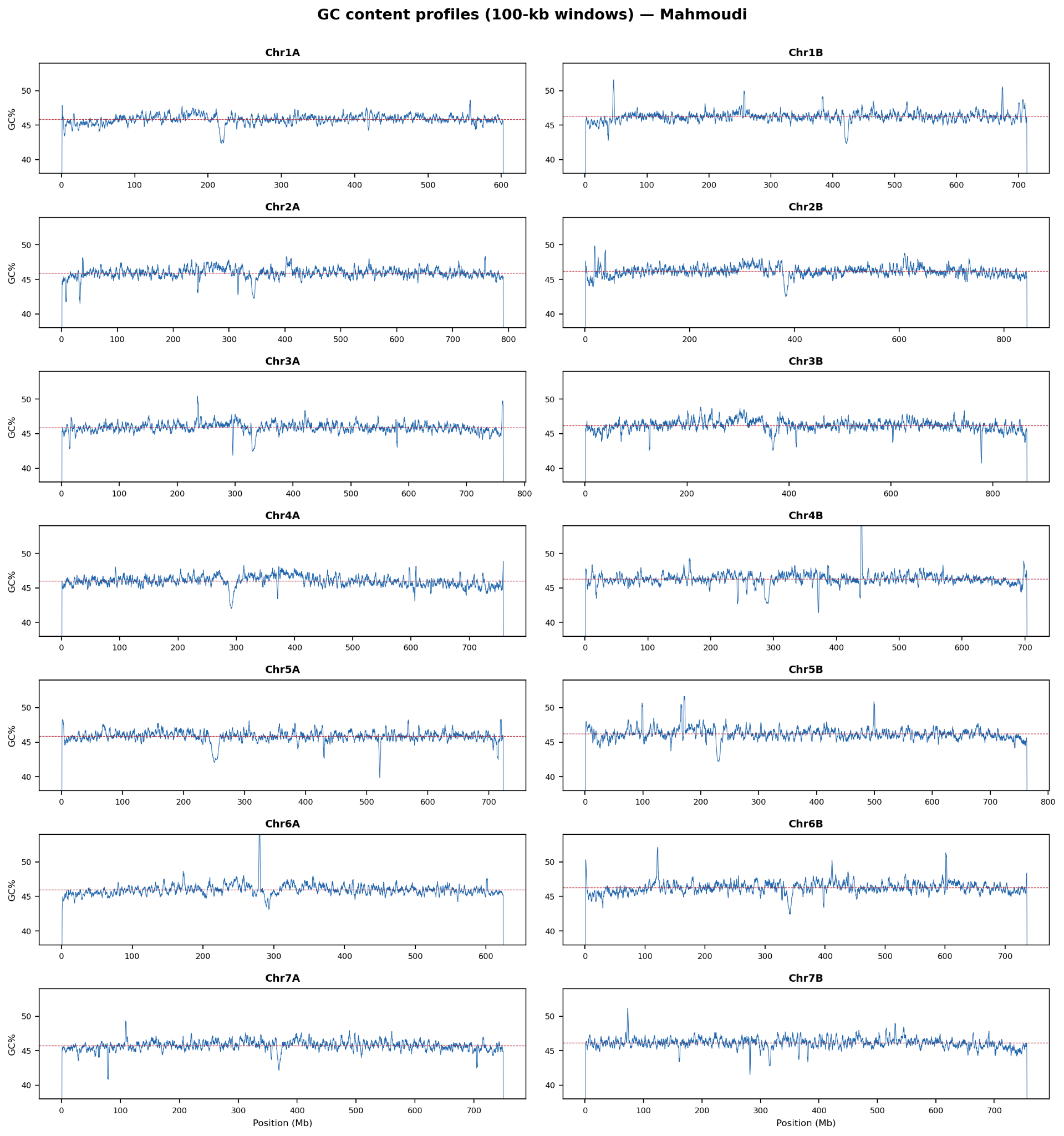

**Figure S6.** GC% calculated in non-overlapping 100 kb windows along each of the 14 chromosomes for Chili (green) and Mahmoudi (blue). GC valleys correspond to centromeric regions; GC peaks correspond to gene-rich chromosome arms. Genome-wide mean GC = 46.08 % (Chili) / 46.03 % (Mahmoudi).

**Supplementary Figure S7. GC skew (G-C)/(G+C) profiles**

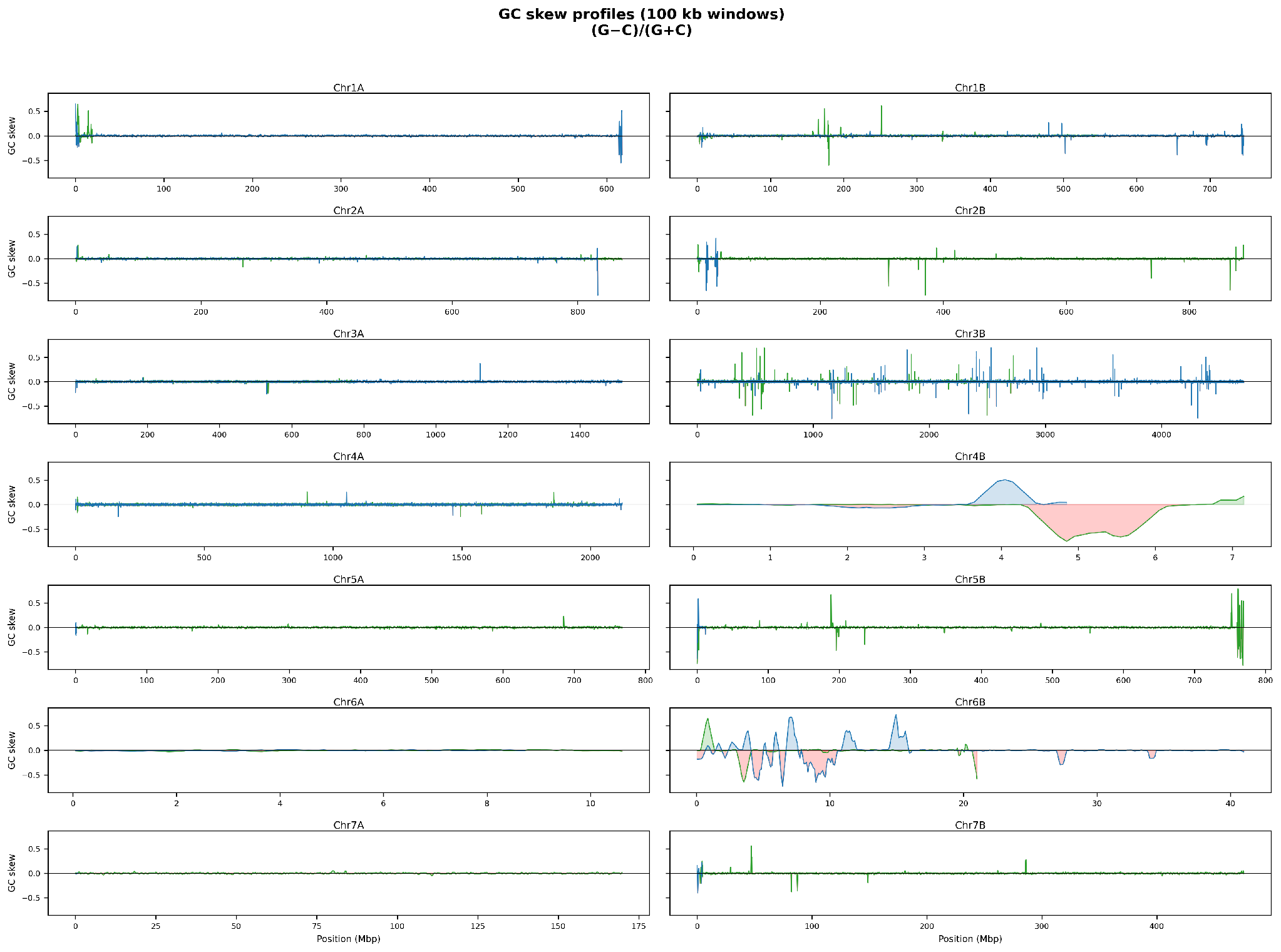

**Figure S7.** Smoothed GC skew per 100 kb window along each chromosome for Chili and Mahmoudi. Positive skew is shaded green and negative skew red; zero-crossings indicate candidate replication-origin / termini positions.

**Supplementary Figure S8. Isochore classification per chromosome**

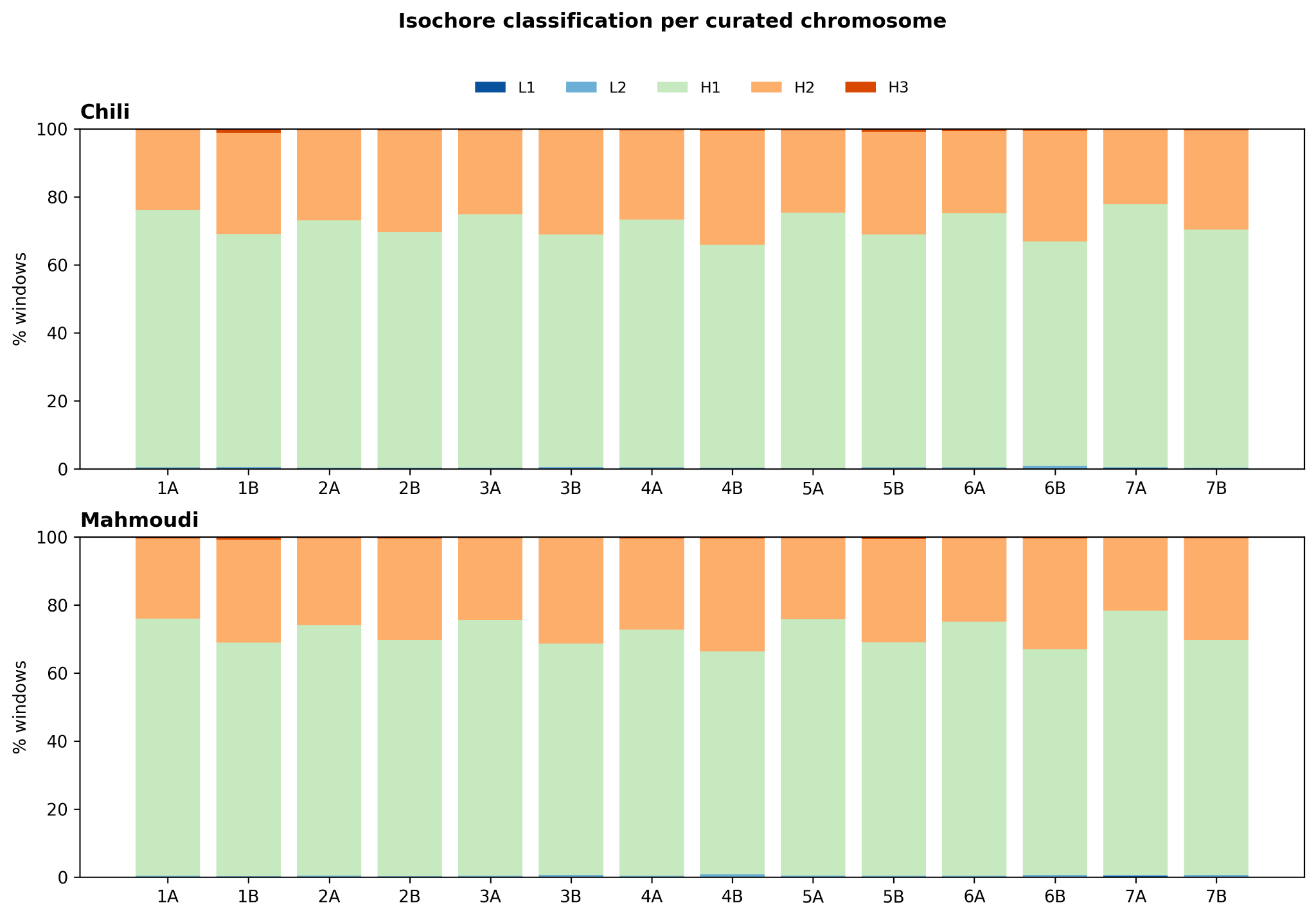

**Figure S8.** Proportion of 100 kb windows assigned to each of five isochore classes (L1 < 37 %, L2 37-41 %, H1 41-47 %, H2 47-52 %, H3 > 52 % GC) for Chili and Mahmoudi. The H1 class dominates (~70.8 % of all windows); the GC-rich H3 class is rare (0.62 % Chili / 0.57 % Mahmoudi).

**Supplementary Figure S9. Predicted centromere positions (GC-valley method)**

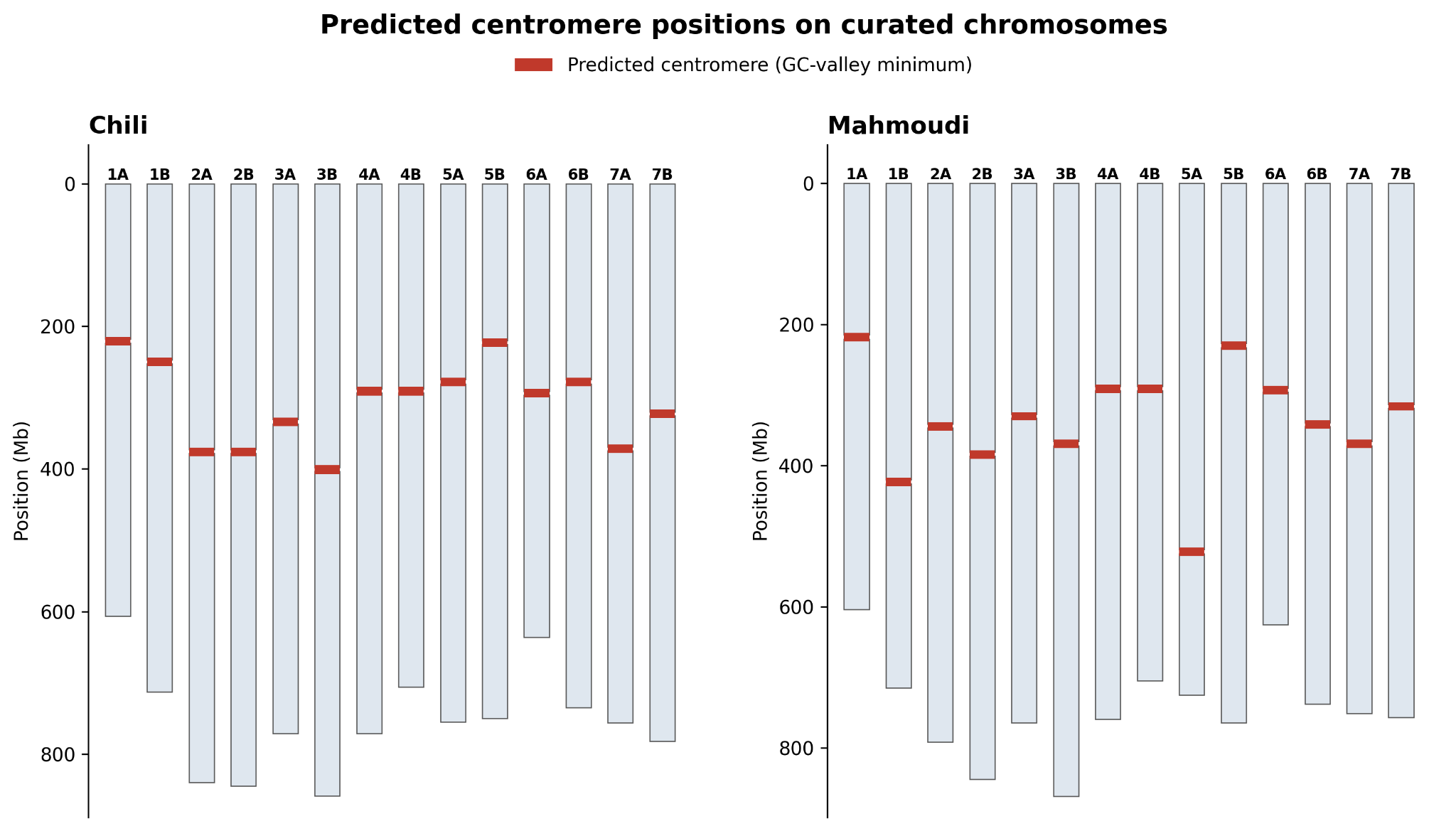

**Figure S9.** Smoothed GC content profiles (11-window moving average over 100 kb windows) for each chromosome of Chili and Mahmoudi. The global GC minimum within the central 60 % of each chromosome is marked by a dashed line and reported as the predicted centromere position.

**Supplementary Figure S10. Ribosomal DNA cluster distribution**

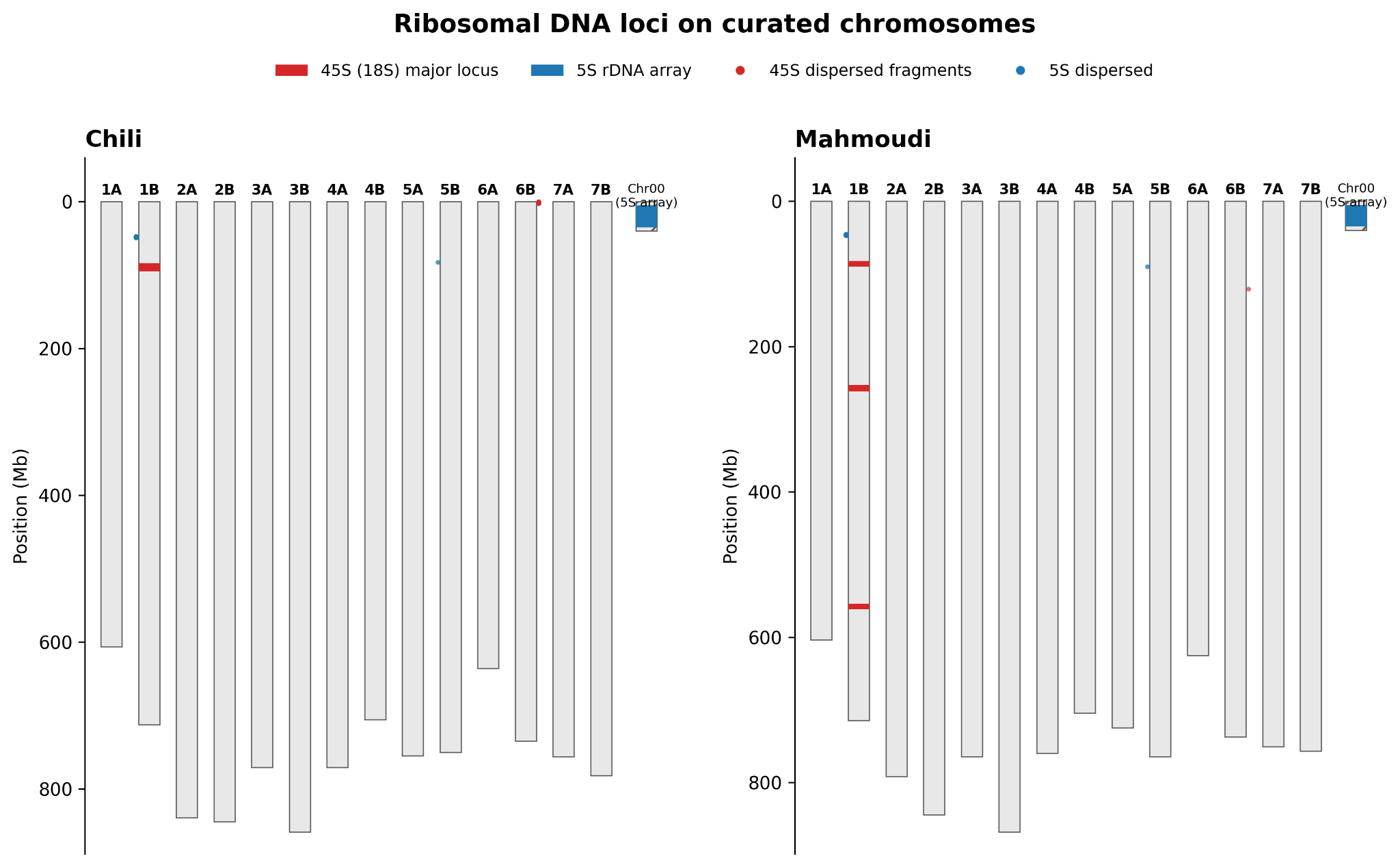

**Figure S10.** Ribosomal DNA loci on the curated chromosomes: principal 45S (18S) locus on Chr1B in both landraces; principal 5S array on an unplaced scaffold (Chr00); dispersed 45S/5S signals.

**Supplementary Figure S11. Dinucleotide composition heatmap (obs/exp ratios)**

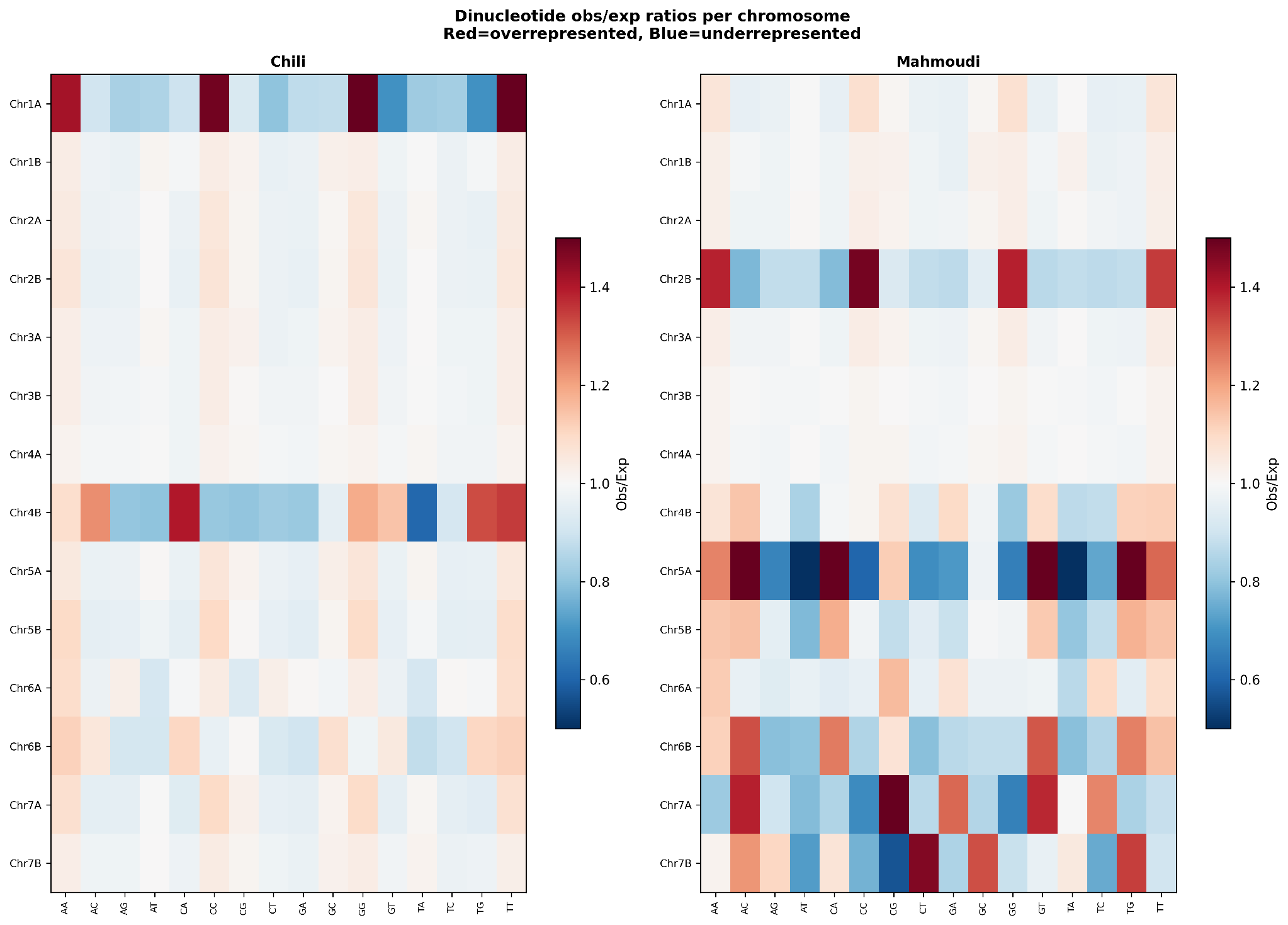

**Figure S11.** Heatmap of observed/expected ratios for all 16 dinucleotides on each chromosome, separately for Chili and Mahmoudi. Red = over-represented (o/e > 1); blue = under-represented (o/e < 1). CpG obs/exp ratios are near 1.0 (0.986 Chili, 1.029 Mahmoudi), consistent with grass genomes (CpG suppression is less pronounced than in mammals).

**Supplementary Figure S12. Alignment-identity distribution vs Svevo v2**

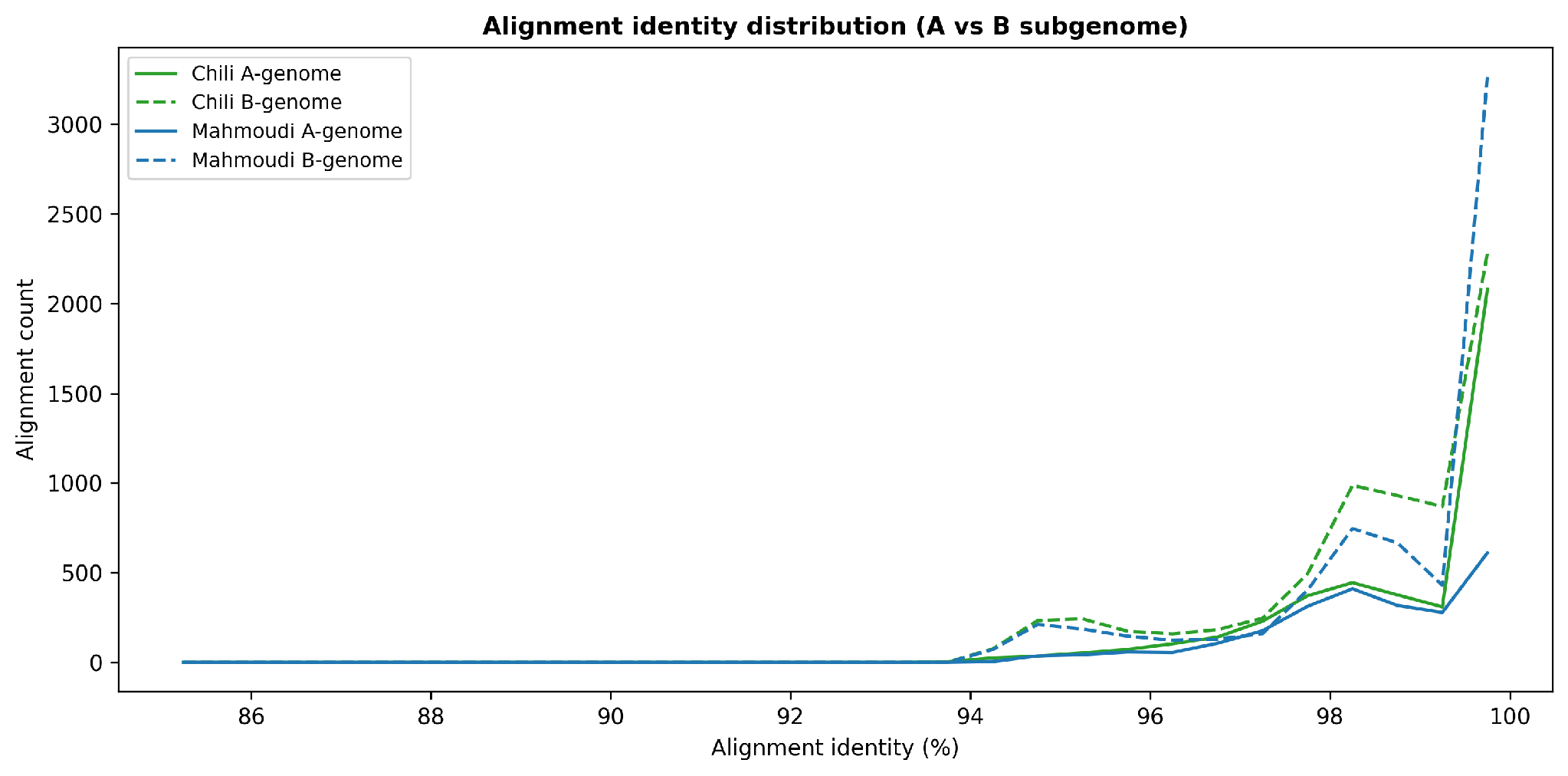

**Figure S12.** Counts of wfmash alignments binned by % identity for the A-subgenome (solid lines) and B-subgenome (dashed); Chili in green, Mahmoudi in blue. Both subgenomes show a bimodal distribution: a peak at 95-98 % (repetitive regions) and a peak at 98-99.5 % (conserved gene-rich regions). Weighted-mean identity = 98.57 % (Chili) / 98.61 % (Mahmoudi).

**Supplementary Figure S13. K-mer sharing heatmap (Chili vs Mahmoudi)**

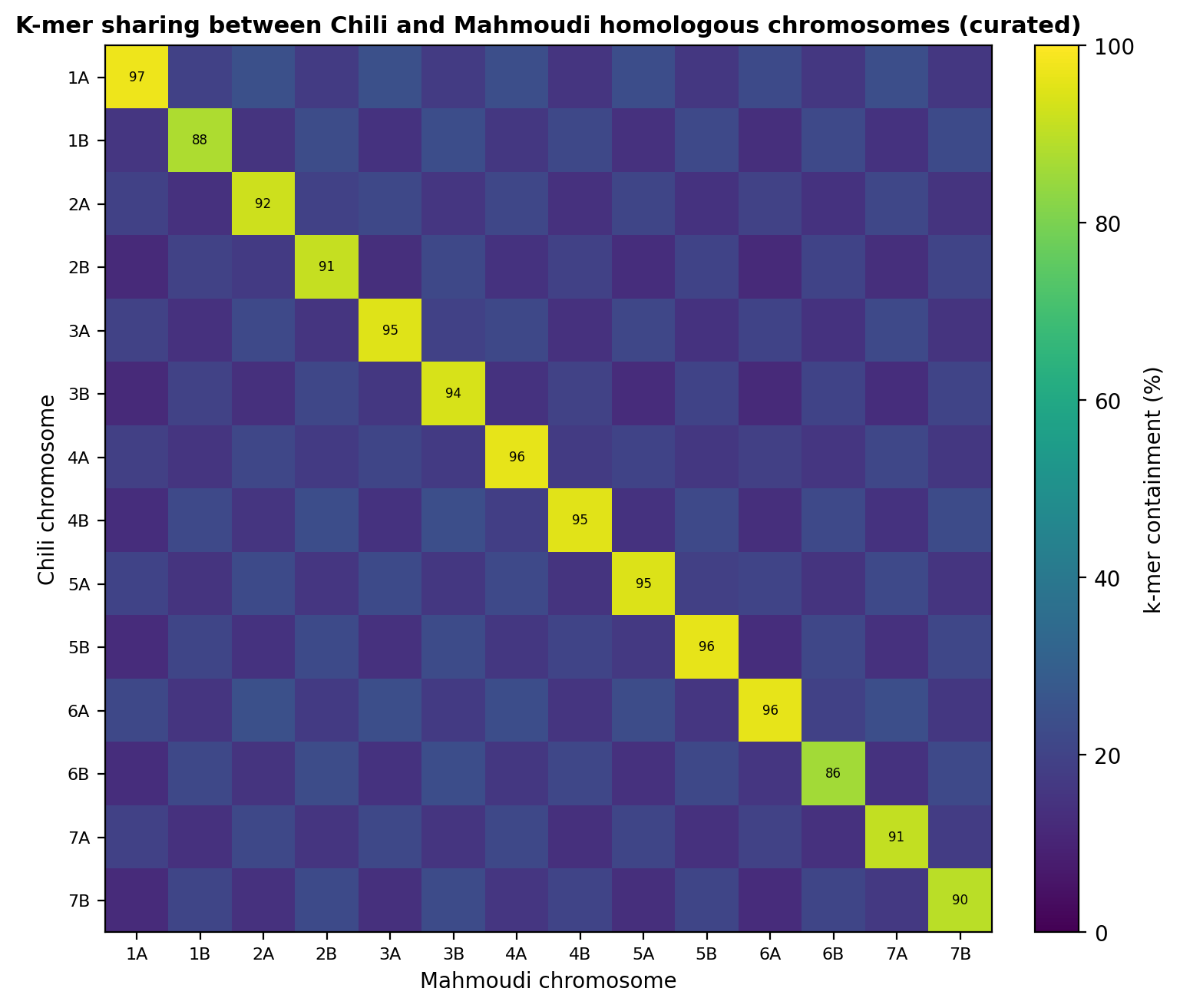

**Figure S13.** Chromosome-level k-mer sharing computed with sourmash containment (k = 21, scaled = 1000). For each of the 14 chromosomes, the heatmap shows the percentage of k-mers shared between Chili and Mahmoudi, the Chili-specific fraction and the Mahmoudi-specific fraction. Mean sharing = 47.3 %, ranging from 93.1 % on the well-assembled Chr2A to 12.3 % on the fragmentary Chr4B.
